# Olfactomedin-4^+^ neutrophils exacerbate intestinal epithelial damage and worsen host survival after *Clostridioides difficile* infection

**DOI:** 10.1101/2023.08.21.553751

**Authors:** A. Huber, S. Jose, A. Kassam, K. N. Weghorn, Maggie Powers-Fletcher, D. Sharma, A. Mukherjee, A. Mathew, N. Kulkarni, S. Chandramouli, M. N. Alder, R. Madan

## Abstract

Neutrophils are key first responders to *Clostridioides difficile* infection (CDI). Excessive tissue and blood neutrophils are associated with worse histopathology and adverse outcomes, however their functional role during CDI remains poorly defined. Utilizing intestinal epithelial cell (IEC)-neutrophil co-cultures and a pre-clinical animal model of CDI, we show that neutrophils exacerbate *C. difficile*-induced IEC injury. We utilized cutting-edge single-cell transcriptomics to illuminate neutrophil subtypes and biological pathways that could exacerbate CDI-associated IEC damage. As such, we have established the first transcriptomics atlas of bone marrow (BM), blood, and colonic neutrophils after CDI. We found that CDI altered the developmental trajectory of BM and blood neutrophils towards populations that exhibit gene signatures associated with pro-inflammatory responses and neutrophil-mediated tissue damage. Similarly, the transcriptomic signature of colonic neutrophils was consistent with hyper-inflammatory and highly differentiated cells that had amplified expression of cytokine-mediated signaling and degranulation priming genes. One of the top 10 variable features in colonic neutrophils was the gene for neutrophil glycoprotein, Olfactomedin 4 (OLFM4). CDI enhanced OLFM4 mRNA and protein expression in neutrophils, and OLFM4^+^ cells aggregated to areas of severe IEC damage. Compared to uninfected controls, both humans and mice with CDI had higher concentrations of circulating OLFM4; and in mice, OLFM4 deficiency resulted in faster recovery and better survival after infection. Collectively, these studies provide novel insights into neutrophil-mediated pathology after CDI and highlight the pathogenic role of OLFM4^+^ neutrophils in regulating CDI-induced IEC damage.

**One Sentence Summary:** Utilizing single-cell transcriptomics, IEC-epithelial co-cultures, and pre-clinical models of CDI, we have identified a subset of neutrophils that are marked by OLFM4 expression as pathogenic determinants of IEC barrier damage after CDI.

## Introduction

*Clostridioides difficile* infection (CDI) is the most common healthcare-associated infection in the US (*1, 2*). *C. difficile* bacteria secrete multiple toxins that are cytotoxic to intestinal epithelial cells (IECs). The compromised gut barrier permits translocation of luminal microbes into submucosa, culminating in recruitment of immune cells that are activated by both microbiota- and host-derived molecules. Although *C. difficile* toxins are the main drivers of initial IEC damage, neutrophils are the critical determinants of clinical outcomes, such that excessive colonic neutrophilia in acute CDI and persistence of neutrophils in colonic lumen later on are associated with prolonged diarrhea and worse disease (*3–8*). Neutrophils are among the first cells to arrive at the site of CDI (*3, 9, 10*), where they release a variety of intracellular products e.g., glycoproteins, reactive oxygen species (ROS), proteases, anti-microbial peptides etc. While neutrophil-derived molecules are released to facilitate pathogen control, they also have the potential to exacerbate host tissue damage due to non-specific nature of their activity.

Despite a key role of neutrophils in regulating CDI outcomes, phenotypic heterogeneity of these cells or the effect of different neutrophil populations in regulating *C. difficile*-associated IEC injury has not been studied. In recent years, the dogma that neutrophils are short-lived, homogenous cells has been challenged, and existence of multiple neutrophil populations at homeostasis and after infection have been identified (*11–20*). Traditionally, neutrophil populations were defined based on cell surface receptor expression, release of effector molecules, nuclear morphology, and maturation status (*21–26*). More recently, single-cell RNA sequencing (scRNAseq) has provided a novel, cutting-edge method of defining neutrophil heterogeneity (*14–17, 27*). Here, we utilized unbiased scRNAseq to determine CDI-induced neutrophil heterogeneity in the bone marrow (BM), blood and colon. Transcriptomics profiling revealed that *Olfactomedin-4* (*Olfm4*), a glycoprotein present in neutrophil specific granules, was upregulated in multiple BM and blood neutrophil populations after CDI. OLFM4 was also among the top 10 variable features in colonic neutrophils isolated from the infected host. Interestingly, OLFM4^+^ neutrophils were enriched for expression of genes associated with degranulation, and higher serum OLFM4 was detected in both humans and mice with CDI, compared to uninfected controls. Further, our exciting data reveal that OLFM4^+^ neutrophils exacerbate CDI-induced epithelial injury, while OLFM4 deficiency improves survival of mice with CDI. Altogether, this study provides the first transcriptomic atlas of CDI-induced neutrophils across different tissues, and we have identified a specific neutrophil population that exacerbates *C. difficile*-induced IEC damage.

## Results

### Neutrophils aggravate *C. difficile* toxin-induced epithelial injury

In CDI, excessive colonic neutrophilia correlates with worse histopathology and adverse clinical outcomes (*3, 10, 28, 29*). But studies that directly examine the impact of neutrophils in regulating CDI-associated IEC damage are lacking. To address this knowledge gap, we co-cultured BM neutrophils with an epithelial cell line (CMT-93) in the presence or absence of *C. difficile* toxins (Fig. 1a). Compared to untreated controls, incubation of IECs with *C. difficile* toxins decreased trans-epithelial resistance (TEER), but addition of BM neutrophils did not alter the toxin-induced drop in TEER (Fig. 1a). After IEC barrier is disrupted, lamina propria (LP) neutrophils are exposed to gut bacteria and their products. To recapitulate this *in vivo* scenario, we used a gram-negative bacterial cell wall product lipopolysaccharide (LPS) – when IECs were incubated with *C. difficile* toxins and BM neutrophils in the presence of LPS, there was a significantly greater decline in TEER (Fig. 1a). These data suggest that LPS, by activating naïve BM neutrophils, exacerbates *C. difficile* toxin-induced IEC damage.

**Figure 1:**
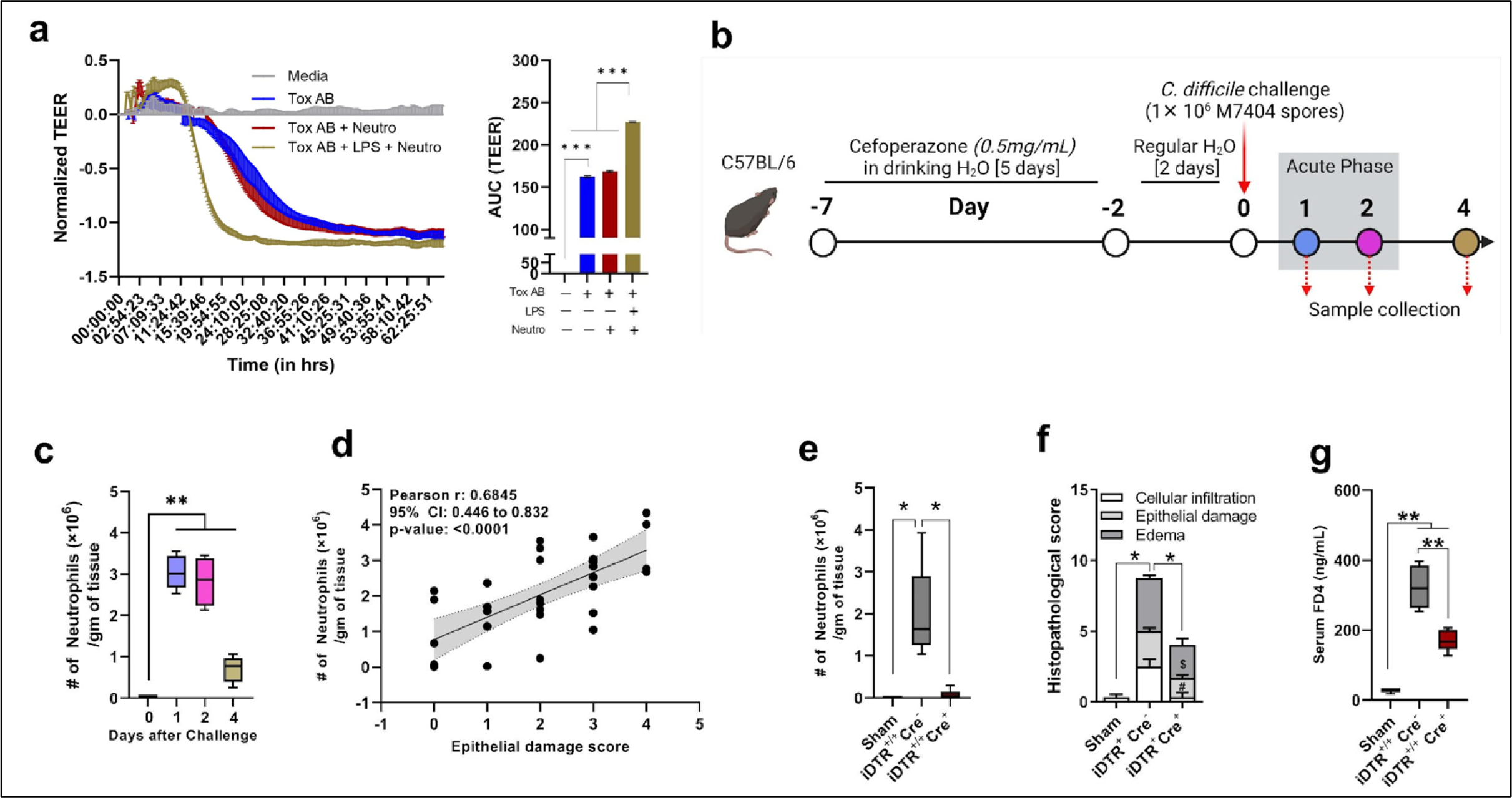
Neutrophils aggravate *C. difficile* toxin-induced epithelial damage. **a)** Change in transepithelial electrical resistance (TEER) of CMT-93 cell monolayers incubated with clostridial toxins A and B, LPS, and neutrophils and area under curve of the TEER plots. Curves are plotted as the normalized mean value of three replicates; representative of 2 independent experiments. **b)** Schematic representation of *C. difficile* challenge experimental design. Age- and gender-matched WT C57BL/6 mice were pre-treated with antibiotics for 5 days in drinking water and challenged with 1×10^6^ *C. difficile* (M7404) spores by oro-gastric gavage 2 days after cessation of antibiotics. Animals were euthanized on days 0, 1, 2, and 4, and samples were collected for further analysis. **c)** Number of neutrophils in colonic tissue of uninfected (day 0) and *C. difficile* infected (day 1, 2 and 4) mice. **d)** Pearson correlation analysis of colonic neutrophil counts and epithelial damage score on day 1 after CDI (N = 33). **e)** Number of neutrophils per gram colonic tissue, **f)** histopathology score and **g)** serum FITC-dextran concentration of uninfected (i.e., sham-challenged) or *C. difficile* infected iDTR^+/+^Cre^-^ and iDTR^+/+^Cre^+^ mice on day 1 after CDI. Panel a is representative of 2 independent experiments, each with triplicate wells; ***p < 0.001, Multiple t-test. For panels c and e-g, data shown as mean ± SEM; N = 3-6 per group; representative of 2 independent experiments (except panel f, 1 experiment); *p < 0.05, Student’s t-test.

To investigate these findings *in vivo*, we examined the association between colonic neutrophil numbers and epithelial damage using a well-established mouse model of CDI (*3, 30–33*) (Fig. 1b & Supp Fig. 1a-f). Acute CDI (i.e., on day 1-2 after infection) resulted in colonic neutrophil accumulation followed by decrease in their numbers during recovery phase (i.e., by day 4 after infection) (Fig. 1c). Importantly, the number of neutrophils per gram colonic tissue in acute CDI correlated very well with epithelial damage score, but not with submucosal edema score (Fig. 1d & Supp Fig. 1g). To directly examine the effect of neutrophils on CDI-induced IEC damage, we used transgenic mice that allow for selective and inducible depletion of endogenous neutrophils (*34*). Mice expressing Cre-inducible simian diphtheria toxin (DT) receptor (DTR) (ROSA26-iDTR) were bred to mice with Cre recombinase under control of a neutrophil-specific MRP-8 promoter (MRP8-Cre^+^) to generate iDTR^+/+^Cre^+^ and iDTR^+/+^Cre^-^ mice (Supp Fig. 1h). DT treatment 12hrs prior to and on the day of CDI depleted endogenous blood and colon neutrophils in iDTR^+/+^Cre^+^ mice, such that their colonic neutrophil numbers were similar to that seen in sham-challenged mice, while control, i.e., iDTR^+/+^Cre^-^ mice had significant neutrophilia during acute infection (Fig. 1e & Supp Fig. 1i-j). iDTR^+/+^Cre^+^ mice also had lower total histopathology scores and less colon length shortening compared to iDTR^+/+^Cre^-^ mice (Fig. 1f & Supp Fig. 1k), but *C. difficile* colony forming units (CFUs) and intra-cecal toxin titers were similar in all mice (Supp Fig. 1l-m). Notably, improved histology score in iDTR^+/+^Cre^+^ was primarily driven by 2 parameters: fewer infiltrating cells and lower epithelial damage score, while submucosal edema score was similar in the two groups (Fig. 1f). Since endogenous neutrophils are depleted in iDTR^+/+^Cre^+^ mice, reduction in cellular infiltration score is expected, but lower CDI-induced epithelial damage score in the absence of neutrophils suggests that these cells are exacerbating IEC barrier injury. The impact of neutrophils on IEC barrier function was examined by a FITC-dextran (FD4) permeability assay: 18 hours after CDI, DT-treated iDTR^+/+^Cre^+^ and iDTR^+/+^Cre^-^ mice were gavaged with FD4 and then plasma samples were collected 6 hours later. Amount of FD4 fluorescence detected in plasma was lower in iDTR^+/+^Cre^+^ mice compared to iDTR^+/+^Cre^-^ (Fig. 1g). Together, these data point to a pathogenic role for neutrophils in acute CDI, such that these cells exacerbate IEC damage caused by *C. difficile* toxins and bacteria, without affecting the pathogen itself.

### Transcriptional atlas of BM and blood neutrophils at homeostasis and in acute CDI

In acute CDI, the number of neutrophils in blood and colon increased while those in BM decreased (Supp Fig. 2a), suggesting migration from BM to blood and to the infected colon. We, therefore, examined the transcriptomic profile of neutrophils from BM, blood, and colon after CDI (Fig. 2a). Since number of neutrophils in intestinal tissue of uninfected mice is extremely low (Fig. 1c and Supp Fig. 2a), it has not been feasible to obtain sufficient cells for single-cell transcriptomics from the colon of uninfected mice (*17*). Thus, from the uninfected control mouse, only BM and blood neutrophils were sequenced. After rigorous quality control (Supp Fig. 3a-e), we obtained high-quality sequences from a total of 32,052 neutrophils: 8656 BM and 4676 blood neutrophils from the uninfected mouse; 7219 BM, 10500 blood and 1001 colonic neutrophils from the infected mouse (Fig. 2b). Unsupervised clustering of sequences obtained from BM and blood neutrophils of uninfected and *C. difficile*-infected mice revealed 9 distinct populations; 6 of which (Neu1-6) were primarily found in BM and 3 (Neu7-9) in blood (Fig. 2c-d & Supp Fig. 3f-l). A combination of unsupervised (i.e., differential gene expression [DGE] and gene ontology [GO] enrichment) and supervised (i.e., scoring using published gene lists associated with cell cycle phases and granule protein expression) analyses revealed that the neutrophil clusters identified in our dataset exhibited distinct gene expression signatures (Fig. 2e-h, Supp Fig. 4a-b, & Supp Tables 1-3). Comparing the cell cycle scores, transcription factor (TF) expression, and granule scores to published studies (*14–16, 27*), we identified Neu1 as pre-neutrophils and Neu2/Neu3 as populations of immature neutrophils (Fig. 2e-g). Based on increased expression of *Cebpb* and decreased expression of specific and gelatinase granule protein genes, Neu4-6 were identified as mature BM neutrophils (Fig. 2e-g), and among these, Neu4 were enriched for genes associated with myeloid leukocyte activation and metabolic processes related to ROS generation (Fig. 2e & h). Neu5 had high amounts of genes related to cell migration, whereas Neu6 had heightened expression of genes contributing to immune responses to bacteria (Fig. 2e & h). Neu7-9 are mature peripheral blood neutrophils enriched for genes associated with cytokine-mediated signaling (Fig. 2e & g-h). Neu8 distinctly expressed genes related to regulation of platelet activation, and Neu9 expressed genes associated with a type I IFN response (Fig. 2e) (*14*). In addition to these independent analyses, we validated neutrophil clusters by comparing our data to previously published neutrophil scRNAseq datasets (*14, 20, 27, 35*). Our findings demonstrated a robust correlation with published reports; Neu1 exhibited similarity to previously defined pre-neutrophils, Neu2-3 were similar to immature neutrophil populations, and Neu4-9 aligned very well with the reported mature neutrophil populations (Supp Fig. 4c-d) (*14, 20, 27, 35*). Leveraging our findings of neutrophil maturation status from each cluster, we performed a pseudotime analysis with Monocle (*36–38*). Neutrophil maturation and differentiation progress in chronological order from Neu1 to Neu9 (Fig 2i & Supp Fig. 4e). To examine the effect of maturation status and aging on driving cellular diversity in our dataset, we correlated pseudotime with maturation gene and aging scores among different populations (Fig. 2j) (*19, 39*). Although maturation scores correlated strongly with pseudotime in the overall dataset (r = 0.84), in Neu5-9, the neutrophil maturation scores plateaued (r = -0.11) (Fig. 2j). Only a weak correlation (r = 0.16) was observed between neutrophil aging score and pseudotime (Fig. 2j). In conclusion, we have defined 9 distinct populations that range from BM pre-neutrophils to mature peripheral blood neutrophils, and our data indicate that maturation status was an important driver of cell diversity. Interestingly, the effect of maturation status was most prominent in early stages of neutrophil development, and factors other than maturation likely contribute to cellular heterogeneity after the Neu5 stage.

**Figure 2:**
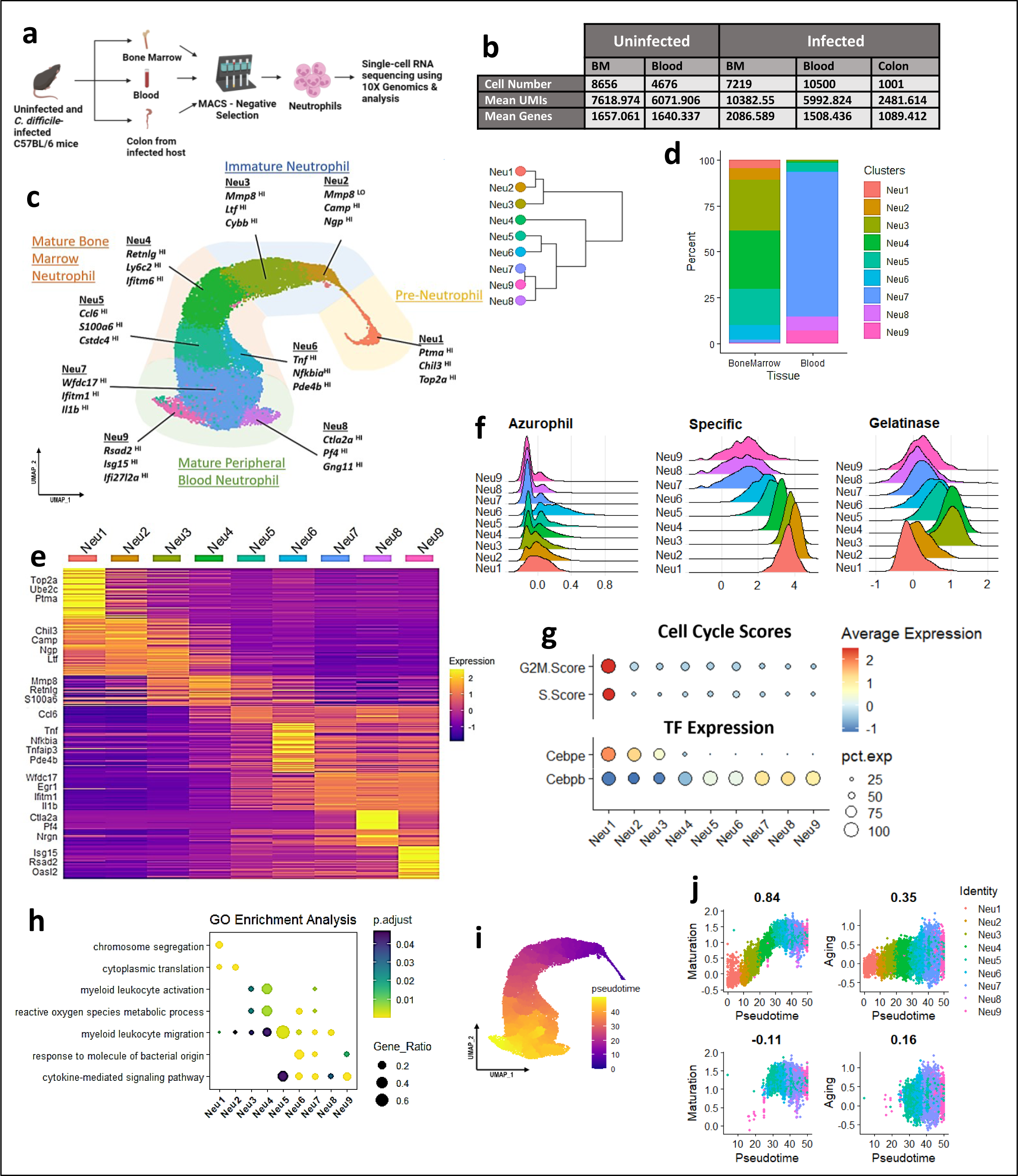
Single-cell transcriptomics reveals neutrophil heterogeneity after CDI. **a)** Bone marrow (BM), blood, and colon were harvested from uninfected and *C. difficile*-infected mice (colon samples from infected host only). MACS-negative selection was utilized to enrich samples for neutrophils, which were subsequently sequenced using 10X Genomics droplet-based sequencing. **b)** Table displaying cell numbers, mean genes and mean unique molecular identifiers (UMIs) per cell for each sample. **c)** Uniform Manifold Approximation and Projection (UMAP) of BM and blood neutrophil populations of uninfected and infected hosts colored by cluster (each point on the UMAP represents a single cell) with a dendrogram relating clusters based on gene expression similarities. Signature genes are listed under each cluster identity. Sections of UMAP are colored by neutrophil maturation (i.e., tan = pre-neutrophils, blue = immature neutrophils, red = mature BM neutrophils, and green = peripheral blood mature neutrophils). **d)** Percent distribution of neutrophil clusters in BM and blood. **e)** Row-scaled average expression of top 50 differentially expressed genes (DEGs) with signature genes listed on the side. **f)** Ridge plots of azurophil, specific, and gelatinase granule protein gene expression scores. **g)** Dot plot of cell cycle scores (G2 to M phase progression and the S phase) and gene expression of transcription factors associated with neutrophil maturation, *Cebpe* and *Cebpb.* Size of dot corresponds to percent of cells in the cluster and color corresponds to average gene expression level. **h)** Dot plot of gene ontology analysis of neutrophil cluster DEGs. Size of dot corresponds to gene ratio (i.e., number of genes associated with the GO term / total number of input genes) and the color corresponds to Benjamini–Hochberg-corrected p-values. **i)** UMAP of BM and blood clusters colored by pseudotime. **j)** Feature scatter plots of pseudotime plotted against maturation and aging scores for total and mature neutrophil populations (Neu5-Neu9). Each point in the scatter plot represents a cell (colored by cluster). Pearson correlations between the two features are displayed above the plots.

### CDI directs mature BM neutrophils towards a pro-inflammatory lineage

To elucidate potential biological mechanisms by which neutrophils in a *C. difficile*-infected host exacerbate IEC damage, we analyzed infection-induced differences in transcriptomes. In BM, CDI resulted in a marked expansion of Neu1 (3.4% → 6.5%), Neu2 (4.8% → 8.1%), and Neu6 (0.5% → 16.7%) with a concurrent decrease of Neu5 (25.8 % → 12.1%) (Fig. 3a-b). In blood, the proportion of Neu5 and Neu7 (0.3% → 6.8%, and 68.5% → 83.3%, respectively) increased, while that of Neu8 and Neu9 (20.2% → 1.9%, and 9.9% → 5.8%, respectively) decreased (Fig. 3a-b). Additionally, transcriptional reprogramming was observed in mature neutrophil populations (Neu6-9) – hierarchal clustering based on average gene expression showed that Neu6-9 formed a separate branch after CDI, but Neu1-5 were very similar to each other both before and after infection (Supp Fig. 5a). Although neutrophils were mostly confined to the same tissue compartment after infection, we observed a CDI-induced shift of approximately 50% of Neu5 cells into blood (Supp Fig. 5b).

**Figure 3:**
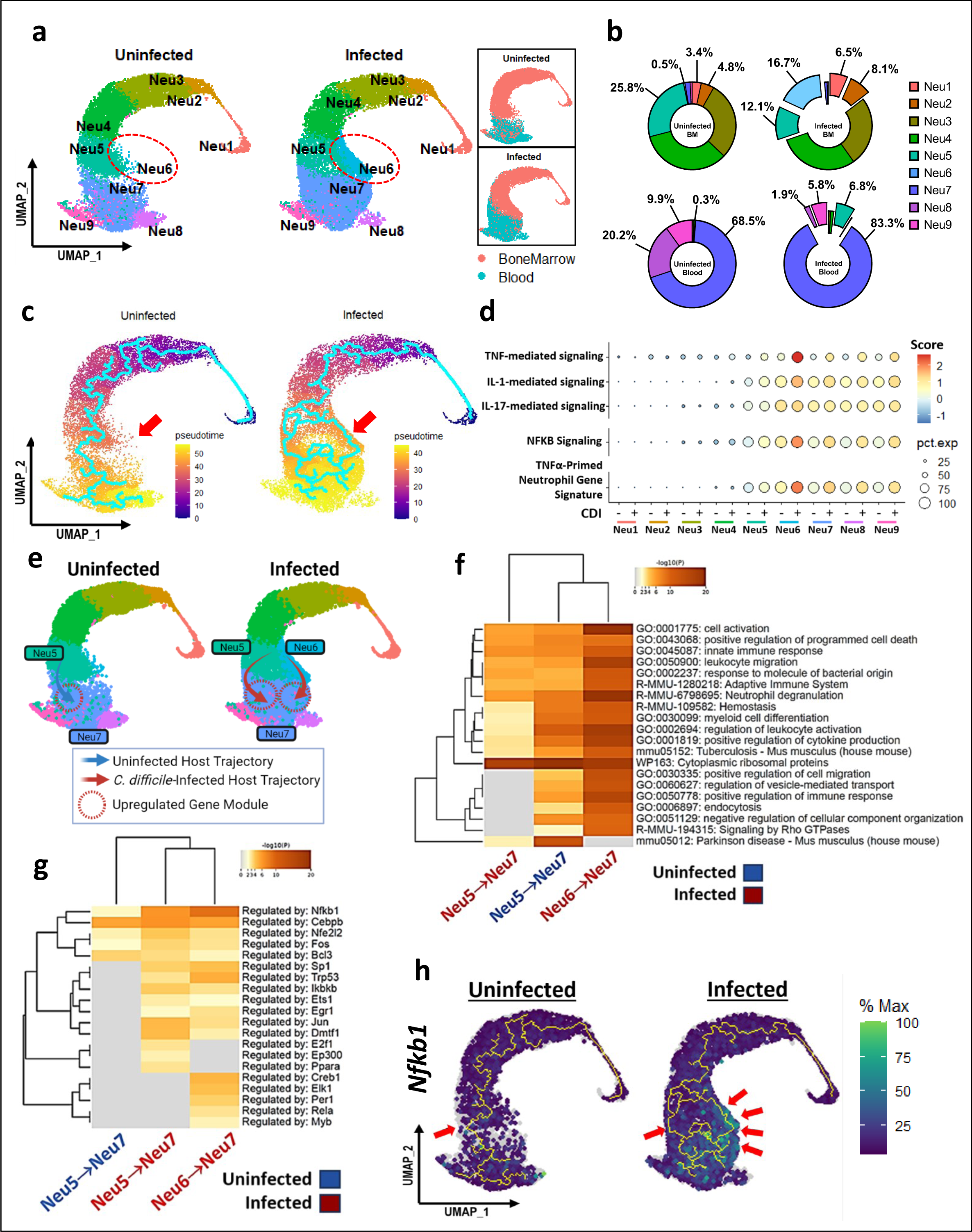
CDI redirects neutrophil differentiation programs to a pro-inflammatory lineage. **a)** UMAP of BM and blood neutrophils of the uninfected and infected hosts colored by clusters and by tissue. **b)** Percent distributions of neutrophil populations by infection status and tissue compartment. **c)** Pseudotime trajectories of neutrophil populations of uninfected versus *C. difficile*-infected hosts. **d)** Dot plot of neutrophil populations of uninfected and infected hosts scored for cytokine-mediated signaling (TNF, IL1, and IL17), downstream mediator pathways (NF-κB), and DEGs of TNFα primed neutrophils. Size of dot corresponds to percent of cells in the cluster and color corresponds to expression level of score. **e)** Monocle branch analysis was utilized to calculate gene modules that change as a function of pseudotime along trajectories that give rise to Neu7. Co-expressed genes upregulated at the end of the trajectories (Neu7) compared to the start of the trajectories (Neu5/Neu6) were used as input for **f)** pathway enrichment analysis plotted in a heatmap. Heatmap cells are colored by p-values and blank squares represent a lack of enrichment for that term for the corresponding cluster. **g)** Predicted transcription factors associated with Neu5/Neu6 to Neu7 differentiation were calculated using TRRUST (Transcriptional Regulatory Relationships Unraveled by Sentence-based Text mining) database. **h)** *Nfkb1* gene expression plotted along pseudotime trajectory in the uninfected and *C. difficile*-infected host. Cells are colored by percent max expression level.

Monocle pseudotime analysis revealed that neutrophil differentiation in an uninfected host proceeds as a single trajectory from Neu1 to Neu7, with Neu7 arising directly from Neu5, and very few Neu6 cells (Fig. 3b-c). After CDI, there was a dramatic increase in the proportion of Neu6 (∼33-fold) and neutrophil differentiation trajectory bifurcated at the level of Neu5, such that there is an additional path of Neu7 development, directly from Neu6 (Fig. 3b-c). Cytokines prime neutrophils to enhance their functions and pro-inflammatory capacity (*40*). Since there was a dramatic infection-induced increase in Neu6, we examined the possibility that these cells develop in response to CDI-induced pro-inflammatory cytokines. GO analysis indeed found high expression of genes that are associated with cytokine-mediated signaling in Neu6 (Fig. 2h & Supp Fig. 4b). Scoring neutrophil clusters using gene lists associated with specific cytokines that are increased in CDI, i.e., TNF, IL-1 and IL-17 (*41–43*), showed that signaling mediated by TNF and its downstream effector NF-κB, was most strongly induced in Neu6 (Fig. 3d). To determine if the transcriptome of Neu6 cells was consistent with that of published TNFα-primed neutrophils, we scored our populations for DEGs specific to these cells (*41*), and found that peak expression of these genes occurred in Neu6 of the infected host (Fig. 3d). Interestingly, while Neu6 highly expressed TNF-induced gene signatures after CDI compared to Neu5, both can differentiate into Neu7 (Fig. 3c). We took two complementary approaches to investigate the impact of TNFα priming on Neu7: unsupervised clustering and DGE analysis of Neu7 (Supp Fig. 5c-e & Supp Table 4), and pathway enrichment analysis on gene modules that change as a function of pseudotime along trajectories of Neu7 differentiation from Neu5/Neu6 to Neu7 (Fig. 3e-f, Supp Fig. 5f, & Supp Tables 5-6). We observed intra-cluster heterogeneity in Neu7 with segregation into 3 groups: Neu7a, Neu7b and Neu7c (Supp Fig. 5c). Pseudotime projection indicated that Neu7a differentiated from Neu5, Neu7b from Neu6 and Neu7c resulted from the convergence of 2 neutrophil trajectories: Neu5 → Neu7 and Neu6 → Neu7 (Fig. 3e & Supp Fig. 5c). Pathway enrichment analysis of the 3 distinct Neu7 development trajectories, i.e., Neu5 → Neu7 in uninfected mouse, and Neu5 → Neu7 + Neu6 → Neu7 after CDI, revealed that cells that arose from Neu6 (i.e., Neu7b) had higher expression of genes that contribute to cell activation, neutrophil degranulation, leukocyte migration, and cytokine production (Fig. 3f). Further, Neu7b had high expression of pro-inflammatory genes, such as *Ccrl2* and *Pde4b* after CDI (Supp Fig. 5d). Consistent with our previous finding that Neu6 significantly expands in response to CDI (Fig. 3b), we observed a substantial increase in Neu7b cells after infection (Supp Fig. 5e). Transcriptional Regulatory Relationships Unraveled by Sentence-based Text (TRRUST) analyses was used to identify candidate TFs that can regulate these differentiation trajectories. Underscoring our observations that TNFα priming influences Neu6 differentiation, TRRUST analysis also predicted that one of the genes that was highly upregulated in the Neu6 → Neu7 trajectory was *Nfkb1* (Fig. 3g-h). Together, these data support a scenario where CDI alters neutrophil differentiation pathways to result in the dramatic expansion of a particular neutrophil population, Neu6. These cells then independently differentiate towards a population with high inflammatory potential (i.e., Neu7b), under the direction of TNFα-NF-κB axis.

### CDI-induced reprogramming amplifies expression of genes associated with degranulation and neutrophil-associated tissue injury

In comparison to the uninfected host, DGE analyses showed significant transcriptomic alterations after CDI in every neutrophil population (Fig. 4a & Supp Table 7). Pathway enrichment analyses based on these differentially expressed genes (DEGs) revealed that neutrophil clusters in the *C. difficile*-infected host globally exhibited significantly higher expression of genes contributing to pro-inflammatory processes like leukocyte migration, cytokine-mediated signaling, and regulation of TNF (Fig. 4b-c & Supp Tables 8-10). Notably, the highest expression signatures found after CDI were in genes contributing to neutrophil degranulation and inflammatory response (Fig. 4b). Degranulation is a critical mechanism to combat infection where activated neutrophils rapidly release intracellular granule proteins (28, 47). We investigated if CDI impacts genes involved in neutrophil granule synthesis and their release. After infection, all neutrophil populations express higher levels of azurophil and gelatinase granule protein genes, with peak expression of azurophil granule genes occurring at Neu6 stage (Fig. 4d-e). Gene expression of specific granule proteins had a slightly different pattern where these were higher in immature (Neu3) and peripheral blood (Neu7-8) neutrophil populations after infection (Fig. 4f). By using the multi-component NADPH oxidase, neutrophils generate ROS to control infectious agents (*44*). We found that neutrophils in the *C. difficile*-infected mouse had upregulation of genes contributing to ROS and NADPH oxidase generation (Fig. 4g-h). Although these neutrophil-derived products (i.e., granule proteins, ROS, etc.) are directed towards pathogen control, due to the non-specific nature of their action, they can also cause collateral tissue damage (*45–48*). Additionally, we examined gene signatures associated with other cellular processes that are known contributors of neutrophil-associated tissue damage, e.g., pyroptosis and neutrophil extracellular trap release (NET-osis) (*49–52*). Although CDI augmented transcripts associated with pyroptosis in all populations from Neu3 to Neu9, peak expression was observed in 2 specific populations: Neu6, whose numbers were also dramatically increased after infection, and Neu9, that were most differentiated on the pseudotime spectrum (Fig. 4i). Somewhat different from pyroptosis, the NET-associated genes were upregulated in all neutrophils (Fig. 4j). Thus, our transcriptomics data suggest that CDI augments BM and blood neutrophil functionality and primes them for release of potentially harmful intra-cellular contents.

**Figure 4:**
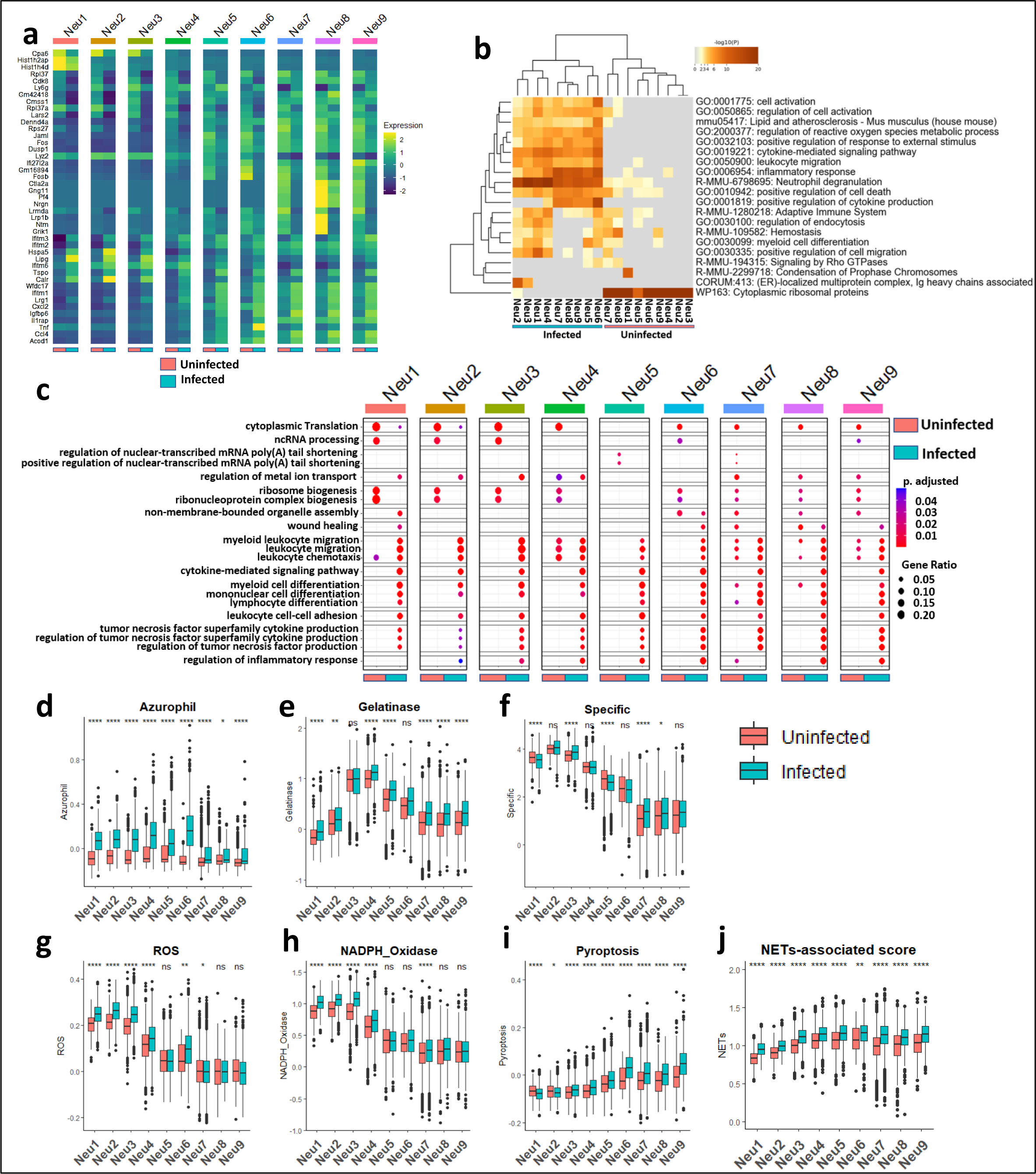
CDI reprograms neutrophil transcriptional landscape and amplifies gene expression associated with degranulation and neutrophil-associated tissue injury. **a)** Row-scaled average expression of DEGs for each cluster compared to their uninfected/*C. difficile*-infected counterparts. Comparative cluster DEGs were utilized for **b)** Metascape gene enrichment analysis and **c)** clusterProfiler GO analysis. Lists of genes associated with neutrophil functions obtained from published literature and gene ontology databases were utilized to score neutrophil populations on gene expression associated with formation of **d)** azurophil granule proteins, **e)** gelatinase granule proteins, **f)** specific granule proteins, **g)** ROS, and **h)** NADPH Oxidase. Additionally, neutrophils were scored on pathways such as **i)** pyroptosis, and **j)** NET-osis. For box plots, the middle line represents median values of gene expression, the upper and lower hinges of box represent first and third quartile (25^th^ and 75^th^ percentiles), and the upper/lower whiskers extend from the hinge to the largest/smallest value no further than 1.5 * interquartile range (IQR) from the hinge (where IQR is the distance between the first and third quartiles). *p < 0.05; **p < 0.01; ***p < 0.001; ****p < 0.0001; Welch’s t-test.

### Infiltrating colonic neutrophils exhibit a hyperinflammatory phenotype during CDI

We were able to obtain high quality sequencing data from colonic neutrophils of a *C. difficile* infected mouse (Fig. 2b & Supp Fig. 3m-q), and therefore had the unique opportunity to examine neutrophil heterogeneity at tissue level. We clustered colonic neutrophils along with BM and blood neutrophils from *C. difficile*-infected mouse and performed pseudo-temporal ordering (Fig. 5a-b & Supp Fig. 3m-q). Colonic neutrophils were located at the end of neutrophil pseudotime spectrum suggesting that these are the most differentiated subpopulation (Fig. 5b). Compared to BM, expression of genes associated with inflammatory response, innate immune response, and regulation of cell activation was higher in both peripheral blood and colonic neutrophils (Fig. 5c & Supp Tables 11-13). While expression of genes associated with degranulation was enriched in neutrophils from all tissue compartments, colonic neutrophils specifically had more transcripts of genes associated with cellular responses to stimuli and regulation of response to a biotic stimulus (Fig. 5c). Scoring of neutrophils from infected mouse using gene lists associated with responses to CDI-induced cytokines (*41–43*) showed that compared to BM and blood, colon neutrophils had higher expression of genes related to TNF-, IL-1-, and IL-17-mediated signaling pathways (Fig. 5d). Independent clustering of colonic neutrophils revealed 2 transcriptionally distinct populations – cNeu1 and cNeu2 (Fig. 5e). DGE analyses of these populations showed high levels of *Nfkbia*, *Bcl3a1b* and *Tnfaip3* in cNeu1 and that of *Rsad2*, *Isg15* and *Retnlg* in cNeu2 (Fig. 5f & Supp Table 14). Using colonic neutrophil DEGs for pathway enrichment analysis, we found that cNeu1 were enriched for genes associated with inflammatory response, cytokine-mediated signaling pathway, NF-κB signaling, and response to molecule of bacterial origin (Fig. 5g & Supp Tables 15-16). cNeu2, on the other hand, were enriched for genes associated with innate immune response and neutrophil degranulation (Fig. 5g). These data suggest that neutrophils migrating to infected colon are highly differentiated and exhibit a hyperinflammatory phenotype with cNeu1 primed for cytokine responses to amplify inflammation and cNeu2 ready for degranulation.

**Figure 5:**
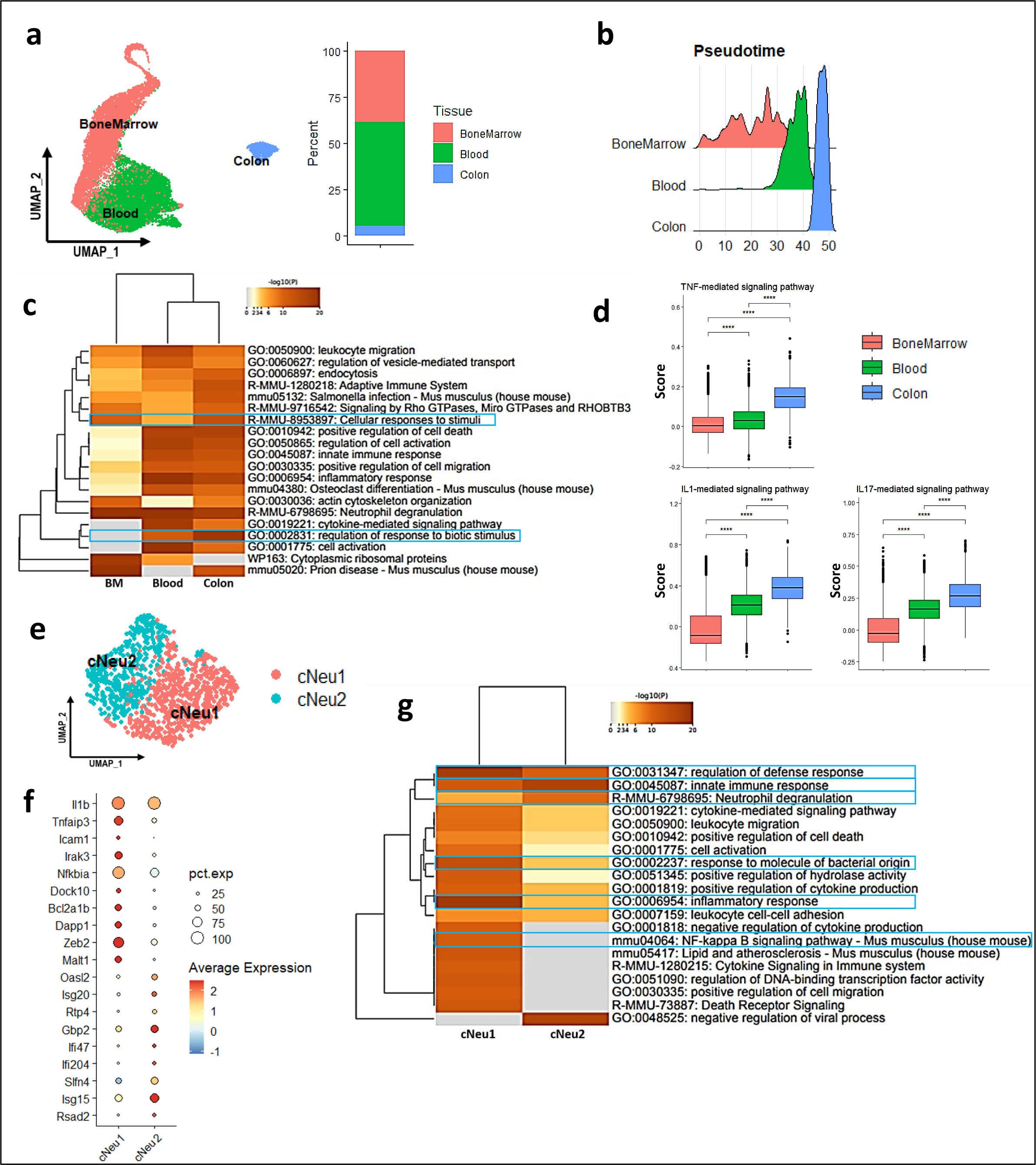
Transcriptional profiling of bone marrow, blood, and infiltrating colon neutrophils in a *C. difficile*-infected host. **a)** UMAP depiction of BM, blood, and colon neutrophils of the *C. difficile*-infected host and percent distributions. **b)** Monocle pseudotime analysis of tissue neutrophils. **c)** Heatmap of pathway enrichment analysis of BM, blood, and colon neutrophils of the infected host. **d)** Boxplots of inflammatory cytokine signature scores (TNF-, IL1-, and IL17-mediated signaling pathways). **e)** UMAP depiction of infiltrating colon neutrophils, and **f)** dot plot of DGE analysis for corresponding clusters. Genes are plotted by scaled average expression. **g)** Heatmap of pathway enrichment analysis of colon neutrophil clusters. *p < 0.05; **p < 0.01; ***p < 0.001; ****p < 0.0001; Kruskal-Wallis test.

### CDI induces neutrophil OLFM4 expression and augments its extra-cellular release

To identify novel neutrophil-associated markers that define heterogeneity in CDI, we plotted the standardized variance against average expression of all transcripts from neutrophils of infected mouse (Fig. 6a). Olfactomedin-4 (*Olfm4*), a glycoprotein that is part of neutrophil specific granules had the highest cell-to-cell variation across all populations (Fig. 6a) and in colonic neutrophils, it was one of the top 10 variable features (Fig. 6b). OLFM4 expression changes based on neutrophil developmental stage such that its mRNA peaks in myelocytes and metamyelocytes with a decrease in more mature populations (*26*). Consistent with published literature, we observed more *Olfm4* transcripts early in the neutrophil pseudotime trajectory (Fig. 6c-d). Although CDI increased OLFM4 expression in multiple populations, Neu3 had most *Olfm4* both before and after CDI (Fig. 6d). Flow cytometry confirmed the presence of OLFM4 protein in BM, blood, and colon neutrophils (Fig. 6e-h). While the number of OLFM4^+^ neutrophils in BM and blood remained unchanged after CDI, their numbers in colonic LP significantly increased (Fig. 6f-h). OLFM4 is released from neutrophils in response to noxious insults (*53, 54*). Pathway enrichment analysis of scRNAseq data revealed that *Olfm4^+^* neutrophils had highest association with pathways related to neutrophil degranulation (Fig. 6i & Supp Tables 17-21), and in fact, *C. difficile* infected mice had higher serum OLFM4 compared to uninfected controls (Fig. 6j). Comparison of OLFM4 staining in BM, blood, and LP neutrophils between uninfected and *C. difficile* infected mice supports the phenomenon of CDI-induced degranulation and OLFM4 extra-cellular release: while the proportion of neutrophils that express OLFM4 in BM and blood was unchanged (Fig. 6k-l), their percentage in LP of infected mice was decreased (Fig. 6m). Mean fluorescent intensity (MFI) of OLFM4 expression showed a similar trend where OLFM4 MFI was not changed in BM and blood neutrophils (Fig. 6n-o), but it decreased in LP neutrophils (Fig. 6p). Thus, our transcriptomics and flow cytometry data suggest that CDI primes OLFM4^+^ neutrophils for degranulation, and this results in OLFM4 release in blood and tissues.

**Figure 6:**
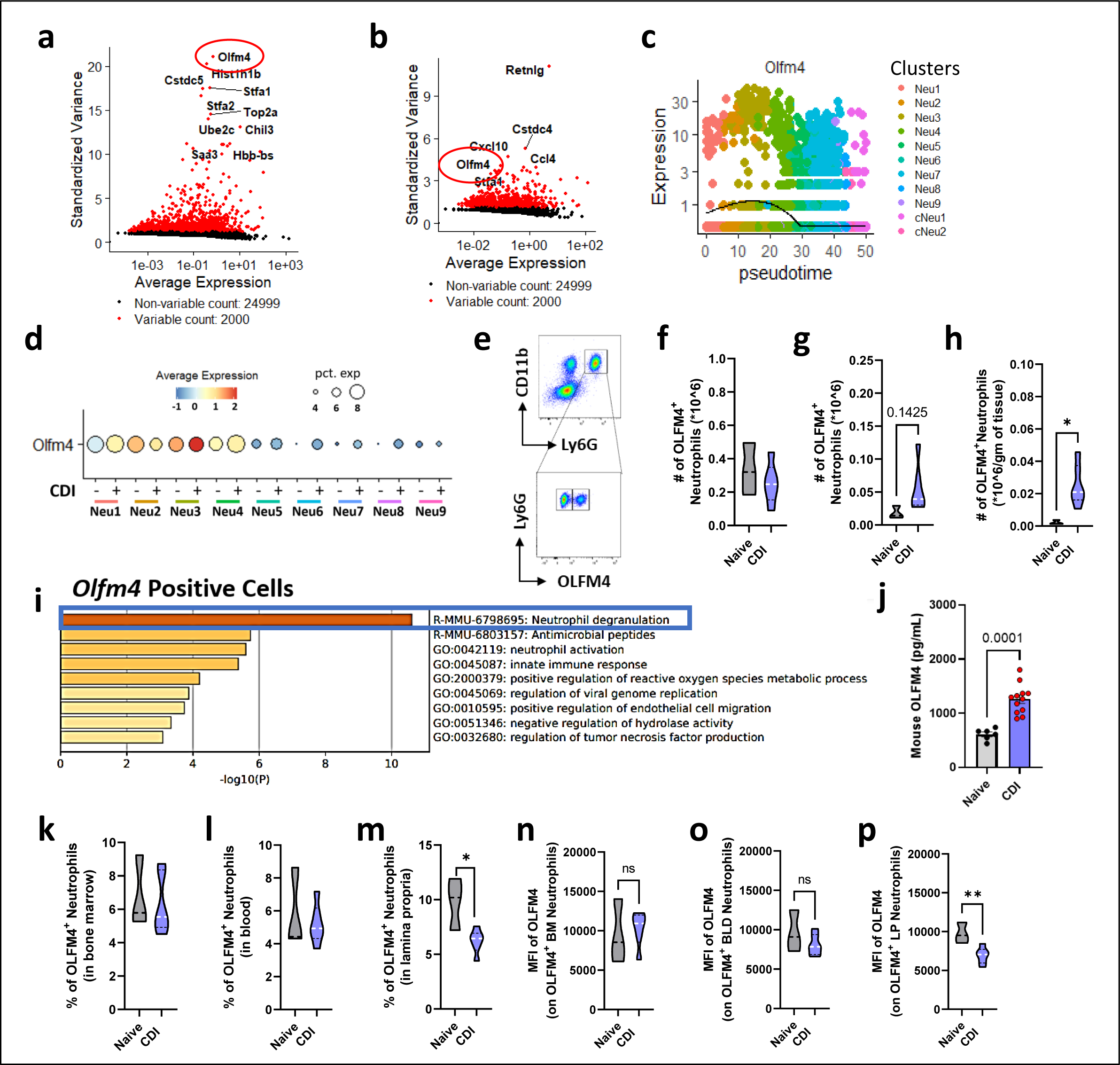
CDI induces OLFM4 expression in neutrophils, and OLFM4^+^ neutrophils are primed for degranulation. **a)** Variability plot of infected host bone marrow, blood, and colon. Genes are plotted by average expression vs standard variance, and the red color represents highly variable genes, while black color represents non-variable genes. **b)** Variability plot of infected host colon. **c)** *Olfm4* dynamics as a function of pseudotime. **d)** Dot plot of *Olfm4* gene expression in uninfected and *C. difficile*-infected hosts. **e)** Representative FACS plot with the gating strategy and number of OLFM4^+^ neutrophils in colonic tissue of naïve and *C. difficile-* infected C57BL/6 mice. Number of OLFM4^+^ neutrophils in **f)** BM, **g)** blood, and **h)** lamina propria of naïve and *C. difficile*-infected mice. **i)** Metascape pathway enrichment analysis of *Olfm4*^+^ neutrophils from the infected host. **j)** Serum OLFM4 levels in naïve and *C. difficile*-infected mice on day 1 after challenge. **k-m)** Percentage of OLFM4^+^ neutrophils and **n-p)** mean fluorescence intensity of OLFM4 in BM (**k and n**), blood (**l and o)**, and lamina propria neutrophils (**l and p)**. Data for panels f-h and j-p shown as mean ± SEM; N = 3-6 per group for all panels except j where N = 6-12; *p < 0.05, **p < 0.01, ***p < 0.001; Student’s t-test.

### OLFM4^+^ neutrophils exacerbate epithelial injury

OLFM4 plays a key role in innate immune responses to infectious agents, and published literature reveal that high proportion of OLFM4^+^ neutrophils in systemic circulation is clearly associated with severe end-organ damage and worse outcomes (*54–62*). Thus, we directly tested the effect of OLFM4 expressing neutrophils in regulating epithelial injury in CDI. To obtain high purity, BM neutrophils from WT and OLFM4 deficient (OLFM4^-/-^) mice were FACS sorted prior to co-culture with CMT-93 cells on a TEER assay plate. Consistent with our prior observation (Fig. 1a), toxins reduced the TEER, but addition of naïve neutrophils (either WT or OLFM4^-/-^) did not change the TEER any further (Fig. 7a-b). As expected, LPS addition enhanced the toxin-induced drop in TEER in the presence of WT neutrophils, but this effect was significantly lower with OLFM4^-/-^ neutrophils (Fig. 7a-b). *In vivo* studies also support a disease enhancing role for OLFM4^+^ neutrophils and point to an adverse impact of these cells on IECs. In WT mice infected with *C. difficile*, OLFM4 staining was more pronounced in areas of worse IEC damage (Fig. 7c; blue arrowheads), compared to those with less damage (Fig. 7c; black arrowheads). Similarly, mice with epithelial damage score in the severe range had more OLFM4^+^ cells per high power field than those with milder injury (Fig. 7d). Interestingly, negligible OLFM4 staining was observed in cecal sections of neutrophil-deficient (iDTR^+/+^Cre^+^) mice compared to neutrophil-sufficient (iDTR^+/+^Cre^-^) mice (Fig. 7e), suggesting that neutrophils are the main source of OLFM4 in colonic tissue after CDI. Taken together, these data imply that OLFM4^+^ neutrophils are clearly pathogenic in nature and they also aggregate to areas of IEC injury.

**Figure 7:**
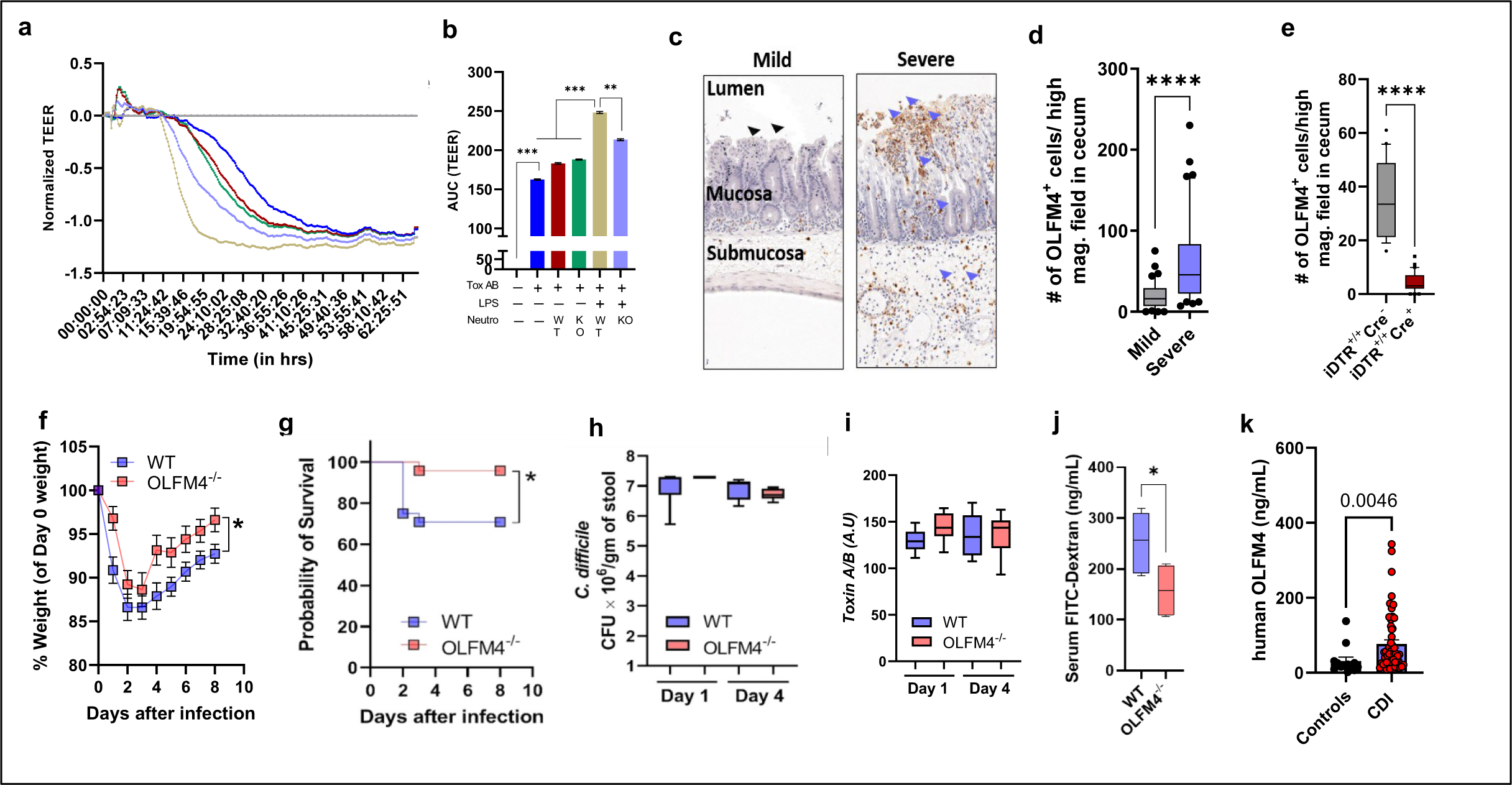
OLFM4^+^ neutrophils exacerbate *C. difficile* toxin-induced epithelial injury, and its deficiency improves host survival after CDI without any effect on the pathogen. **a)** Dynamic measurement of transepithelial electrical resistance (TEER) of CMT-93 cells treated with *C. difficile* toxins, LPS, and BM neutrophils isolated from WT and OLFM4^-/-^ mice. **b)** Area under the curve of the TEER plot. Curves are plotted as the normalized mean value of three replicates; representative of 2 independent experiments. **c)** Representative images of immunohistochemical staining for OLFM4 in areas of low vs. high epithelial damage in cecal sections of *C. difficile*-infected mice. Number of OLFM4^+^ cells per high magnification field in cecal sections **d)** WT mice, and **e)** iDTR^+/+^Cre^-^ (control) iDTR^+/+^Cre^+^ mice, one day after *C. difficile* challenge; N = 6 mice per group; 30-40 fields counted; ****p < 0.001; Student’s t-test. **f)** Percent weight change (compared to day 0 weight) after CDI in mice challenged with M7404 spores. **g)** Difference in survival rate of WT and OLFM4^-/-^ mice after CDI. *C. difficile* h) bacterial colony forming unit (CFU), and **i)** toxin levels per gram of cecal content of *C. difficile*-challenged WT and OLFM4^-/-^ mice on day 1 and day 4 after challenge. **j)** Serum FITC-dextran concentration in *C. difficile*-challenged WT and OLFM4^-/-^ mice on day 4 after infection. Data shown as mean ± SEM; N = 5-6; representative of 2-3 independent experiments for all panels f and h-j; *p < 0.05; Multiple t-test (for f) and Student’s t-test (for h-j). For panel g, N = 15-18; Log-rank [Mantel-Cox] test; *p < 0.05. **k)** Serum OLFM4 levels in patients with and without CDI. Data shown as mean ± SEM; N = 13 controls and 53 *C. difficile*-infected patients; Student’s t-test.

### OLFM4 deficiency improves host survival after CDI without any effect on pathogen

To further examine OLFM4’s role in CDI pathogenesis, we infected age- and gender-matched WT and OLFM4^-/-^ mice with *C. difficile* spores (M7404; 10^6^ spores) (Fig. 1b). The onset of diarrhea, duration of diarrhea and clinical score (assigned based on coat ruffling, posture, ocular discharge and activity level) were comparable between the two groups (Supp Fig. 6a-b). However, OLFM4^-/-^ mice had improved weight gain and a significant survival advantage compared to WT mice (Fig. 7f-g). Impact of OLFM4 on disease severity was more pronounced after infection with a high toxin producing *C. difficile* strain (VPI 10463) where all parameters, i.e., diarrhea severity, clinical score, weight loss and CDI-induced mortality were significantly better in OLFM4^-/-^ mice (Supp Fig. 6c-g). Of note, OLFM4 deficiency did not affect the amount of *C. difficile* (Fig. 7h) or toxin titers (Fig. 7i) in intra-cecal contents. Histologic scoring of H&E-stained cecal sections from *C. difficile* (M7404) infected WT and OLFM4^-/-^ mice showed that there was no difference in the histology scores on day 1 after infection, but by day 4, OLFM4^-/-^ mice had improved scores compared to WT mice (Supp Fig. 6h). Specifically, epithelial damage score was better in OLFM4^-/-^ mice, while mucosal edema and cellular infiltration was similar between the 2 groups (Supp Fig. 6i-k). Consistent with reduced epithelial damage, amount of FD4 detected in plasma samples of OLFM4^-/-^ mice was significantly lower than WT mice, on day 4 after CDI (Fig. 7j). To determine the relevance of OLFM4 in humans, we quantified serum OLFM4 concentration in patients with acute CDI and individuals with diarrhea who tested negative for *C. difficile*. Similar to our murine model, CDI patients had significantly higher systemic OLFM4 compared to controls (Fig. 7k).

## Discussion

In patients with CDI, white blood cell count (which is predominated by neutrophils) of >15x10^9^/L is commonly used to predict disease severity (*63*), and a high number of colonic tissue neutrophils is associated with worse CDI outcomes (*64*). Despite correlative evidence for many years suggesting a critical role for neutrophils in CDI pathogenesis, direct experimental evidence has been lacking. Our studies reveal for the first time that neutrophils exacerbate CDI by worsening IEC barrier integrity. Interestingly, the harmful effect of neutrophils was uncovered only in the presence of pathogenic stimuli from both gram-positive (i.e., *C. difficile* toxins) and gram-negative (i.e., LPS) bacteria. The initial toxin-induced damage to IECs in CDI can lead to dissemination of luminal microbes to deeper tissues, thereby exposing neutrophils to various bacteria. Thus, our findings highlight a key role for combination of bacterial virulence factors in orchestrating a neutrophil activation program that aggravate IEC injury. In contrast, other reports have shown that depleting neutrophils in mice with CDI resulted in increased mortality and uncontrolled bacterial dissemination (*9*), thus revealing a dichotomous role of neutrophils in CDI pathogenesis. A potential explanation for the duality of neutrophils could be the existence of various neutrophil subtypes with pathogenic vs. beneficial potential that are induced in response to CDI. However, so far, no studies have examined the scope of CDI-induced neutrophil heterogeneity or its effect on CDI outcomes. We addressed this key knowledge gap by performing scRNAseq on neutrophils isolated from multiple tissue compartments of uninfected and *C. difficile*-infected mice. In doing so, we have established the first neutrophil transcriptomics atlas after CDI. While previous studies have reported high-quality transcriptomes from BM and blood neutrophils at homeostasis and after noxious insults, tissue neutrophils have been technically challenging to isolate and examine (*14, 17*). Since CDI is characterized by robust tissue neutrophilia response, this infection model provided a unique opportunity to isolate and characterize colonic neutrophils too. Using cutting-edge 10X Genomics droplet-based platform for sequencing, we were able to obtain high-quality data from 1001 colonic neutrophils.

At homeostasis, BM and blood neutrophils clustered into nine distinct populations ranging from pre-neutrophils to mature neutrophils. While the same populations were present after CDI, their proportions and tissue distribution were significantly altered. The typical host response to an infection includes a rapid egress of neutrophils from BM reservoir into circulation and expansion of precursor populations to meet the increased demand of these cells (*16, 65*). We observed a similar phenomenon after CDI where proportion of mature neutrophils (i.e., Neu5) increased in blood with a concurrent decrease of these cells in BM. At the same time, relative numbers of pre-neutrophils (i.e., Neu1) and immature neutrophils (i.e., Neu2) increased in BM. Thus, we have identified Neu1, Neu2 and Neu5 as key components of CDI-induced expedited generation and extravasation of neutrophils from BM.

An important finding of our studies is the CDI-induced diversion of neutrophil differentiation trajectory towards a pro-inflammatory intermediate, i.e., Neu6. Prior to infection, only a few BM cells could be categorized as Neu6, but CDI increased this population ∼33-fold. Importantly, Neu6 transcriptome was consistent with neutrophils undergoing TNFα-priming. Thus, our data suggests that CDI-induced pro-inflammatory mediators, such as TNFα, divert neutrophil differentiation in the BM (I.e., Neu5) to a priming stage (i.e., Neu6), which results in the differentiation of peripheral blood neutrophils with heightened expression of genes associated with activation, leukocyte migration, and cytokine production. Our findings therefore provide an explanation for how a pathogen that is mostly restricted to the colonic lumen also impacts innate immune cell development and their release at distant locations in BM and blood; and point to TNFα as a vital link that orchestrates neutrophil responses across various tissue compartments after CDI. Interestingly, genes involved in processes of neutrophil-associated tissue injury e.g., granule protein synthesis, ROS production and toxic mediator release, i.e., degranulation, pyroptosis and NET-osis, were all upregulated in BM and blood neutrophils after CDI. Additionally, we present novel findings that are specific to tissue neutrophils in the context of CDI – cells that infiltrate into the infected site exhibit gene signatures that are consistent with augmented response to pro-inflammatory cytokines (i.e., TNF, IL-1, and IL-17), along with expressing genes that affect degranulation. Altogether, our findings support a scenario in which CDI alters the neutrophil differentiation program and enhances effector functions, with neutrophil-mediated tissue damage being a possible side-effect of this augmented functionality.

Leveraging transcriptomics data, we identified *Olfm4* as the most highly variable gene in BM, blood and colon, and one of the top 10 variable features in colon. Prior studies in humans and animal models have identified that circulating OLFM4^+^ neutrophil percentage and its serum level have the potential to predict clinical disease severity after an infectious insult (*61, 66*), and reviewed in detail by Liu and Rodgers (*55*). Our study highlights a similar pathogenic role for these cells in CDI, but additionally we present primary evidence of their role in aggravating the tissue damage that is caused by pathogens, i.e., IEC injury mediated by *C. difficile* and its toxins. In cell cultures, neutrophils from OLFM4^-/-^ mice caused less damage compared to WT neutrophils, and when challenged with *C. difficile,* OLFM4^-/-^ mice exhibited less epithelial damage and improved IEC barrier functions compared to WT mice. OLFM4 is a neutrophil specific granule protein that is released during degranulation and NET-osis (*53*). Neutrophil activation results in rapid release of intracellular granule proteins to fight pathogens (*49, 67*), but high local concentrations of certain neutrophil-derived products can also cause bystander tissue damage (*48*). Compared to *Olfm4^-^* neutrophils, we observed higher expression of genes associated with degranulation in *Olfm4^+^*neutrophils; and at a protein level, OLFM4 MFI was lower in tissue neutrophils of *C. difficile*-infected mice compared to uninfected mice suggesting OLFM4 release after CDI. Since more neutrophil recruitment is expected at areas of severe IEC damage, and OLFM4 staining was also more pronounced in the same area, our data indicate that CDI directs the transcriptional programs of tissue neutrophils to enhance OLFM4 release, potentially via degranulation. OLFM4 secretion upon exposure to a strong secretagogue has been substantiated by prior research (*53*), and we also observed that CDI enhanced OLFM4 levels in the serum of infected mice. Our exciting data from OLFM4^-/-^ mice illustrates two key points about its role in disease pathogenesis. First, OLFM4-deficiency had no effect on *C. difficile* burden or toxin titers, indicating that OLFM4 effects are mainly targeted towards the host. Second, the improved clinical parameters in OLFM4^-/-^ mice were seen only during the recovery phase of infection while acute CDI was similar to WT mice, suggesting that OLFM4 could inhibit pathways related to recovery from CDI, for example, those that contribute to IEC repair and restitution. Our observation of CDI patients having more serum OLFM4 compared to uninfected individuals highlights importance of this molecule in the context of clinical disease. The clinical spectrum of CDI ranges from mild, self-limited diarrheal illness to severe colitis and sometimes death. Notably, the proportion of circulating OLFM4^+^ neutrophils in humans exhibits wide variation (range from 5-50%) (*68*), and these cells clearly exacerbate CDI-associated epithelial damage. Thus, differences in OLFM4^+^ neutrophil percent prior to CDI and in response to infection could influence CDI severity and clinical disease outcomes.

Despite making key observations on neutrophil programming, heterogeneity and its role in driving CDI-associated IEC injury, our studies do have some limitations. The temporal relationship between transcriptional changes, protein accumulation and release of OLFM4 in BM, blood and colonic neutrophil populations remains to be elucidated. OLFM4 protein interacts with frizzled receptors, a vital component of Wnt/β-catenin pathway involved in proliferation and survival of IECs (*69*). Thus, OLFM4 could directly affect the repair of IEC barrier in CDI; but whether OLFM4 augments pathways of cell damage or inhibits IEC repair and regeneration or affects both is unknown and will be a focus of future studies. Although we show that CDI enhances proportion of OLFM4^+^ neutrophils and serum OLFM4 concentration, the direct association of these markers with clinical disease in humans is not addressed. Understanding the relationship between OLFM4 and CDI severity can provide valuable insights into the pathogenesis and prognosis of this disease, and therefore a prospective study to examine the impact of OLFM4^+^ neutrophils percent and its serum concentration, on CDI severity is warranted.

In sum, we provide crucial experimental evidence of a pathogenic role for neutrophils in CDI and detail the first transcriptomics atlas of CDI-induced neutrophils across various tissues. Our data from pre-clinical animal models and a cohort of CDI patients clearly point to OLFM4 as a molecule of interest in understanding the inter-individual variability in CDI severity, and suggest that OLFM4^+^ neutrophils and/or serum OLFM4 levels have the potential to be used as biomarkers to risk stratify CDI patients.

## Materials and Methods

### Mouse Model of CDI

All animals were maintained and bred at the VMU, Cincinnati Veterans Affairs Medical Center, under pathogen-free conditions in individually ventilated cages. In CDI experiments, eight to thirteen-week-old C57BL/6 wildtype (WT) or OLFM4 deficient (OLFM4^-/-^) mice were challenged with purified *C. difficile* spores (1×10^4^ VPI 10463 spores or 1×10^6^ M7404 spores per mouse) by oral gavage and monitored daily as described previously (*30, 70*).

### Neutrophil Isolation and *in vitro* assays

Bone marrow neutrophils were sorted from WT and OLFM4^-/-^ mice using a MACS mouse neutrophil isolation kit (Miltenyi Biotech #130-097-658, Germany) or a Sony MA900 sorter as described previously(*3*). Neutrophil purity and percentage of cells expressing OLFM4 was evaluated by flow cytometry as described elsewhere (*3, 54*). For TEER assays, sorted neutrophils were added to CMT-93 (mouse colonic epithelial cells; ATCC) monolayer on 96-well E-plate (Agilent # 300601010), and changes in TEER were measured every 15 minutes for up to 72hrs as described previously (*71*) and in Supplementary methods.

### Single-Cell RNAseq

Clinical disease, pathogen burden and colonic damage was similar in all 3 infected WT C57Bl/6 mice (Supp Fig. 2b-f). Tissues were collected from the mouse with maximum weight loss on day 2 after infection (indicated with red color in Supp Fig. 2b-d) and neutrophils were enriched using MACS beads-based negative selection prior to sequencing. Although tissue collection and sequencing of cells from the uninfected host was performed at a later data than the *C. difficile*-infected host, we used an age-matched WT C57Bl/6 male mouse from the same colony and sequenced the samples with same protocols and on the same instruments. All samples were collected at ∼10 am (or 4 hours after the lights turn on, Zeitgeber time 4), and processed as previously described (*3, 30, 31, 70*). Neutrophil sorting buffer was supplemented with 1U/µl of RNase inhibitor. For colon samples, in addition to magnetic beads that were part of the kit, mouse CD326 (EpCAM) MicroBeads were also used to reduce the amount of IEC contamination. Cells were then resuspended in media with 1U/µl RNase inhibitor and 10% FCS at room temperature. Cell viability was determined by trypan blue exclusion. All samples sent for sequencing had viability >70%. Sequencing was performed using Chromium X scRNA-seq and Illumina platform equipment at CCHMC Single Cell Genomics Core. For cell preparation, NextGEM assay was used and sequencing with 3’v3.1 assay chemistry.

### Post-Sequencing Filtering and Quality Control

Cell Ranger 7.0.0 was utilized for sequencing read alignment to the mm10 mouse transcriptome and quantification of the expression of cellular transcripts. Output files were analyzed using Seurat 4.0 in which filtering data set for low quality cells, doublets, cells with mitochondrial, hemoglobin, and ribosomal genes, and non-neutrophils was performed. See supplementary methods for more details.

### Annotation and Clustering

Annotation of neutrophils was performed by using: (i) AUCell and neutrophil-specific gene lists from PangloaDB; (ii) SingleR to validate clusters with Tabula Muris Senis dataset; and (iii) ToppGene database (See supplementary methods for more details) (*72–76*). Neutrophils of BM and blood of uninfected and infected hosts were integrated and clustered using Seurat. Cells were clustered using the first 15 principal components as input to perform the RunUMAP function and the resolution of the clusters we decided by using Clustree and BuildCluster tree analyses (*77*). See supplementary methods for more details.

For other analyses including infected host tissue compartment neutrophil analysis and colon neutrophil analyses, these cells were clustered at dimensions 1:12 and 1:10 at resolutions 0.3 and 0.3, respectively. Samples were clustered utilizing Seurat Standard Workflow without integration.

### Differential Gene Expression Analysis

After finalizing neutrophil clusters, we found signature genes using FindAllMarkers (used to find DEGs in both uninfected and infected samples together) and FindConservedMarkers (used to find DEGs separately).. For pathway enrichment analyses, DEGs obtained from FindAllMarkers function were utilized.

### Pathway Enrichment Analyses

To calculate biological processes associated with DEGs, clusterProfiler was utilized to find Gene Ontology (GO) terms with gene ratios and Benjamini–Hochberg-corrected P*-*values (*78*). Pathway enrichment, TF regulator prediction, and gene overlap analyses were performed by inputting gene lists obtained from DGE analysis into Metascape (*79*).

### Pseudotime Analysis

Monocle (version 3) was utilized to infer developmental trajectories of neutrophil populations (*36–38*). A single-cell trajectory was constructed to understand relationships of each cluster in the neutrophil lineage. Neu1 was chosen as the root_cells based off neutrophil scoring showing it was the least mature population (based off TF expression and proliferation scores) and aspects of DEG and gene ontology suggesting it is a pre-neutrophil population. Trajectory analysis was performed in a similar manner for uninfected and infected samples, separately, and for all infected tissues. Additionally, Monocle was used to subset branches of the trajectories and obtain gene modules associated with changes in pseudotime of selected branches.

### Neutrophil Module Scoring

Module scoring was used to calculate the average normalized expression of corresponding gene lists associated with neutrophil biological functions. Lists of genes were obtained from Gene Ontology databases and previous literature (Supp table 22). Cell cycle scoring was done by using CellCycleScoring function in Seurat with genes from Nestorowa et al. (*80*). Statistical differences were calculated with a Welch’s t-test for comparisons of clusters from uninfected and infected hosts and Kruskal-Wallis tests for comparisons of neutrophils of different tissue compartments in the infected host (*p < 0.05; **p < 0.01; ***p < 0.001; ****p < 0.0001).

### Variable Gene Expression Analysis

For analysis of tissues from *C. difficile*-infected mice, we also utilized Seurat’s VariableFeatures function to find features exhibiting high cell-to-cell variation in the dataset. Standard variance vs average expression was then plotted with labels of the top ten variable genes.

### *In vivo* Epithelial permeability assay

Neutrophils in iDTR^+/+^Cre^+^ mice were depleted with intraperitoneal diphtheria toxin (500ng; Sigma-Aldrich) injection and challenged with *C. difficile* spores, as described previously (*32, 34*). The mice were then gavaged with 100uL of 25mg/mL FITC Dextran (FD4; Sigma) at 18h after spore challenge, and amount of FD4 in plasma was estimated using fluorescence spectrophotometry. Littermate iDTR^+/+^Cre^-^ mice were used as controls. In experiments to evaluate the effect of OLFM4 on epithelial barrier integrity after CDI, WT, and OLFM4^-/-^ mice were gavaged with FD4 18hrs after *C. difficile* spore challenge, and its plasma concentration was estimated.

### Human data collection and analysis

Discarded plasma samples were collected from hospitalized patients at the University of Cincinnati (UC) Medical Center who had diarrheal illness and whose stool samples were sent for *C. difficile* testing. Samples were collected within 48hrs of *C. difficile* testing and after initial storage at 4°C for a maximum of 24hrs, they were stored at −80°C before further testing. We collected samples from 53 *C. difficile* cases (patients with diarrhea and stool sample positive for *C. difficile* TcdB by PCR) and 13 controls (patients with diarrhea and stool sample negative for *C. difficile* TcdB by PCR). Patient data collection and analysis were approved by the UC Institutional Review Board (IRB); #2019-0195.

### ELISA

Plasma was separated from blood samples by centrifugation at 5,000x g at 4^°^C for 5 minutes and stored at -80^°^C until use. Levels of OLFM4 in plasma were measured using human (abcam; Cat# ab267805) and mouse (LSBio; Cat# LS-F6064-1) OLFM4 ELISA Kits as recommended by the manufacturers.

### Estimation of *C. difficile* pathogen burden and toxin

Pathogen load of *C. difficile* in mice was determined by plating 1:10 dilutions of cecal contents on brain heart infusion supplement (BHIS) bacterial culture plates containing taurocholate. Plates were incubated anaerobically at 37°C for 24h and colonies were counted manually to enumerate colony forming units (CFU) per gram of cecal content. The level of toxins in cecal contents was determined using the *C. difficile* TOXA/B ELISA (TechLab, VA, USA) according to the manufacturer’s instructions.

## Histopathology of cecal tissue sections

Cecal tissue samples were fixed in Bouin’s solution (Sigma) overnight. Samples were washed and dehydrated in 70% ethanol prior to paraffin embedding. Four-micron sections were stained with hematoxylin and eosin (H&E) and scored for inflammatory cell infiltration, edema, and epithelial disruption. A score of 0 to 4, denoting increasingly severe abnormality, was assigned for each of these parameters by a pathologist in a blinded fashion.

## Statistical analysis

All statistical analyses were performed using GraphPad Prism 5.0 software (GraphPad software Corporation, Inc, CA, USA). For comparison of groups, a student’s t-test, or multiple t-tests were used. Differences in survival after CDI were analyzed using log-rank Mantel-cox test. A p-value below 0.05 was considered significant. For gene ontology analyses, Benjamini–Hochberg-corrected p*-*values were used instead of unadjusted p-values. For comparisons between neutrophil clusters from uninfected and infected mice, statistical differences were calculated with a Welch’s t-test (*p < 0.05; **p < 0.01; ***p < 0.001; ****p < 0.0001). For comparisons of tissue neutrophil clusters, statistical differences between groups were calculated by Kruskal-Wallis test (*p < 0.05; **p < 0.01; ***p < 0.001; ****p < 0.0001).

## Acknowledgments

We thank Dr. George S. Deepe Jr. for critical reading of the manuscript and valuable discussions; and Research Flow Cytometry Core and Single-Cell Genomics core at CCHMC for their technical assistance.

## Author contributions

S.J., M.N.A. and R.M. conceptualized and designed the study. A.H. performed scRNA analysis and ELISAs. S.J., A.H., A.M. and A.M. conducted *in vitro* experiments, animal studies and flow cytometry. D.S. reviewed and scored histopathology slides. N.K. and S.C. conducted the pathogen burden experiments. A.K., K.N.W., M.P.F. assisted in patient sample collection and processing. A.H., S.J. and R.M. wrote the manuscript. All authors reviewed the manuscript. R.M. obtained funding for the project.

## References

1. F. C. Lessa, Y. Mu, W. M. Bamberg, Z. G. Beldavs, G. K. Dumyati, J. R. Dunn, M. M. Farley, S. M. Holzbauer, J. I. Meek, E. C. Phipps, L. E. Wilson, L. G. Winston, J. A. Cohen, B. M. Limbago, S. K. Fridkin, D. N. Gerding, L. C. McDonald, Burden of Clostridium difficile infection in the United States. N Engl J Med 372, 825–834 (2015).

2. A. Y. Guh, Y. Mu, L. G. Winston, H. Johnston, D. Olson, M. M. Farley, L. E. Wilson, S. M. Holzbauer, E. C. Phipps, G. K. Dumyati, Z. G. Beldavs, M. A. Kainer, M. Karlsson, D. N. Gerding, L. C. McDonald, Trends in U.S. Burden of Clostridioides difficile Infection and Outcomes. N Engl J Med 382, 1320–1330 (2020).

3. O. Horrigan, S. Jose, A. Mukherjee, D. Sharma, A. Huber, R. Madan, Leptin Receptor q223r Polymorphism Influences Clostridioides difficile Infection-Induced Neutrophil CXCR2 Expression in an Interleukin-1β Dependent Manner. Frontiers in Cellular and Infection Microbiology 11, (2021).

4. K. Solomon, A. J. Martin, C. O’Donoghue, X. Chen, L. Fenelon, S. Fanning, C. P. Kelly, L. Kyne, Mortality in patients with Clostridium difficile infection correlates with host pro-inflammatory and humoral immune responses. J Med Microbiol 62, 1453–1460 (2013).

5. R. E. El Feghaly, J. L. Stauber, E. Deych, C. Gonzalez, P. I. Tarr, D. B. Haslam, Markers of intestinal inflammation, not bacterial burden, correlate with clinical outcomes in Clostridium difficile infection. Clin Infect Dis 56, 1713–1721 (2013).

6. R. E. El Feghaly, J. L. Stauber, P. I. Tarr, D. B. Haslam, Intestinal inflammatory biomarkers and outcome in pediatric Clostridium difficile infections. J Pediatr 163, 1697–1704 e1692 (2013).

7. M. M. Abhyankar, J. Z. Ma, K. W. Scully, A. J. Nafziger, A. L. Frisbee, M. M. Saleh, G. R. Madden, A. R. Hays, M. Poulter, W. A. Petri, Jr., Immune Profiling To Predict Outcome of Clostridioides difficile Infection. mBio 11, (2020).

8. C. C. Lee, J. C. Lee, C. W. Chiu, P. J. Tsai, W. C. Ko, Y. P. Hung, Neutrophil Ratio of White Blood Cells as a Prognostic Predictor of Clostridioides difficile Infection. J Inflamm Res 15, 1943–1951 (2022).

9. I. Jarchum, M. Liu, C. Shi, M. Equinda, E. G. Pamer, Critical role for MyD88-mediated neutrophil recruitment during Clostridium difficile colitis. Infect Immun 80, 2989–2996 (2012).

10. S. Jose, R. Madan, Neutrophil-mediated inflammation in the pathogenesis of Clostridium difficile infections. Anaerobe 41, 85–90 (2016).

11. C. Summers, S. M. Rankin, A. M. Condliffe, N. Singh, A. M. Peters, E. R. Chilvers, Neutrophil kinetics in health and disease. Trends Immunol 31, 318–324 (2010).

12. L. G. Ng, R. Ostuni, A. Hidalgo, Heterogeneity of neutrophils. Nature Reviews Immunology 19, 255-265 (2019).

13. M. Beyrau, J. V. Bodkin, S. Nourshargh, Neutrophil heterogeneity in health and disease: a revitalized avenue in inflammation and immunity. Open Biol 2, 120134 (2012).

14. X. Xie, Q. Shi, P. Wu, X. Zhang, H. Kambara, J. Su, H. Yu, S. Y. Park, R. Guo, Q. Ren, S. Zhang, Y. Xu, L. E. Silberstein, T. Cheng, F. Ma, C. Li, H. R. Luo, Single-cell transcriptome profiling reveals neutrophil heterogeneity in homeostasis and infection. Nat Immunol 21, 1119–1133 (2020).

15. D. E. Muench, A. Olsson, K. Ferchen, G. Pham, R. A. Serafin, S. Chutipongtanate, P. Dwivedi, B. Song, S. Hay, K. Chetal, L. R. Trump-Durbin, J. Mookerjee-Basu, K. Zhang, J. C. Yu, C. Lutzko, K. C. Myers, K. L. Nazor, K. D. Greis, D. J. Kappes, S. S. Way, N. Salomonis, H. L. Grimes, Mouse models of neutropenia reveal progenitor-stage-specific defects. Nature 582, 109–114 (2020).

16. M. Evrard, I. W. H. Kwok, S. Z. Chong, K. W. W. Teng, E. Becht, J. Chen, J. L. Sieow, H. L. Penny, G. C. Ching, S. Devi, J. M. Adrover, J. L. Y. Li, K. H. Liong, L. Tan, Z. Poon, S. Foo, J. W. Chua, I. H. Su, K. Balabanian, F. Bachelerie, S. K. Biswas, A. Larbi, W. Y. K. Hwang, V. Madan, H. P. Koeffler, S. C. Wong, E. W. Newell, A. Hidalgo, F. Ginhoux, L. G. Ng, Developmental Analysis of Bone Marrow Neutrophils Reveals Populations Specialized in Expansion, Trafficking, and Effector Functions. Immunity 48, 364–379.e368 (2018).

17. I. Ballesteros, A. Rubio-Ponce, M. Genua, E. Lusito, I. Kwok, G. Fernández-Calvo, T. E. Khoyratty, E. van Grinsven, S. González-Hernández, J. Nicolás-Ávila, T. Vicanolo, A. Maccataio, A. Benguría, J. L. Li, J. M. Adrover, A. Aroca-Crevillen, J. A. Quintana, S. Martín-Salamanca, F. Mayo, S. Ascher, G. Barbiera, O. Soehnlein, M. Gunzer, F. Ginhoux, F. Sánchez-Cabo, E. Nistal-Villán, C. Schulz, A. Dopazo, C. Reinhardt, I. A. Udalova, L. G. Ng, R. Ostuni, A. Hidalgo, Co-option of Neutrophil Fates by Tissue Environments. Cell 183, 1282–1297.e1218 (2020).

18. M. E. Deerhake, E. Y. Reyes, S. Xu-Vanpala, M. L. Shinohara, Single-Cell Transcriptional Heterogeneity of Neutrophils During Acute Pulmonary Cryptococcus neoformans Infection. Front Immunol 12, 670574 (2021).

19. M. Palomino-Segura, J. Sicilia, I. Ballesteros, A. Hidalgo, Strategies of neutrophil diversification. Nat Immunol 24, 575–584 (2023).

20. Z. Ai, Revealing key regulators of neutrophil function during inflammation by re-analysing single-cell RNA-seq. PLoS One 17, e0276460 (2022).

21. C. Silvestre-Roig, A. Hidalgo, O. Soehnlein, Neutrophil heterogeneity: implications for homeostasis and pathogenesis. Blood 127, 2173–2181 (2016).

22. Y. Tsuda, H. Takahashi, M. Kobayashi, T. Hanafusa, D. N. Herndon, F. Suzuki, Three different neutrophil subsets exhibited in mice with different susceptibilities to infection by methicillin-resistant Staphylococcus aureus. Immunity 21, 215–226 (2004).

23. C. L. Zindl, J. F. Lai, Y. K. Lee, C. L. Maynard, S. N. Harbour, W. Ouyang, D. D. Chaplin, C. T. Weaver, IL-22-producing neutrophils contribute to antimicrobial defense and restitution of colonic epithelial integrity during colitis. Proc Natl Acad Sci U S A 110, 12768–12773 (2013).

24. J. Pillay, B. P. Ramakers, V. M. Kamp, A. L. Loi, S. W. Lam, F. Hietbrink, L. P. Leenen, A. T. Tool, P. Pickkers, L. Koenderman, Functional heterogeneity and differential priming of circulating neutrophils in human experimental endotoxemia. J Leukoc Biol 88, 211–220 (2010).

25. S. Bauer, M. Abdgawad, L. Gunnarsson, M. Segelmark, H. Tapper, T. Hellmark, Proteinase 3 and CD177 are expressed on the plasma membrane of the same subset of neutrophils. J Leukoc Biol 81, 458–464 (2007).

26. S. N. Clemmensen, C. T. Bohr, S. Rorvig, A. Glenthoj, H. Mora-Jensen, E. P. Cramer, L. C. Jacobsen, M. T. Larsen, J. B. Cowland, J. T. Tanassi, N. H. Heegaard, J. D. Wren, A. N. Silahtaroglu, N. Borregaard, Olfactomedin 4 defines a subset of human neutrophils. J Leukoc Biol 91, 495–500 (2012).

27. R. Grieshaber-Bouyer, F. A. Radtke, P. Cunin, G. Stifano, A. Levescot, B. Vijaykumar, N. Nelson-Maney, R. B. Blaustein, P. A. Monach, P. A. Nigrovic, O. Aguilar, R. Allan, J. Astarita, K. F. Austen, N. Barrett, A. Baysoy, C. Benoist, B. D. Brown, M. Buechler, J. Buenrostro, M. A. Casanova, K. Chowdhary, M. Colonna, T. Crowl, T. Deng, F. Desland, M. Dhainaut, J. Ding, C. Dominguez, D. Dwyer, M. Frascoli, S. Gal-Oz, A. Goldrath, T. Johanson, S. Jordan, J. Kang, V. Kapoor, E. Kenigsberg, J. Kim, K. w. Kim, E. Kiner, M. Kronenberg, L. Lanier, C. Laplace, C. Lareau, A. Leader, J. Lee, A. Magen, B. Maier, A. Maslova, D. Mathis, A. McFarland, M. Merad, E. Meunier, P. A. Monach, S. Mostafavi, S. Muller, C. Muus, H. Ner-Gaon, Q. Nguyen, G. Novakovsky, S. Nutt, K. Omilusik, A. Ortiz-Lopez, M. Paynich, V. Peng, M. Potempa, R. Pradhan, S. Quon, R. Ramirez, D. Ramanan, G. Randolph, A. Regev, S. A. Rose, K. Seddu, T. Shay, A. Shemesh, J. Shyer, C. Smilie, N. Spidale, A. Subramanian, K. Sylvia, J. Tellier, S. Turley, B. Vijaykumar, A. Wagers, C. Wang, P. L. Wang, A. Wroblewska, L. Yang, A. Yim, H. Yoshida, C. ImmGen, The neutrotime transcriptional signature defines a single continuum of neutrophils across biological compartments. Nature Communications 12, 2856 (2021).

28. J. C. Brazil, C. A. Parkos, Pathobiology of neutrophil-epithelial interactions. Immunol Rev 273, 94–111 (2016).

29. H. A. Carpenter, N. J. Talley, The importance of clinicopathological correlation in the diagnosis of inflammatory conditions of the colon: histological patterns with clinical implications. Am J Gastroenterol 95, 878–896 (2000).

30. S. Jose, A. Mukherjee, M. M. Abhyankar, L. Leng, R. Bucala, D. Sharma, R. Madan, Neutralization of macrophage migration inhibitory factor improves host survival after Clostridium difficile infection. Anaerobe 53, 56–63 (2018).

31. S. Jose, M. M. Abhyankar, A. Mukherjee, J. Xue, H. Andersen, D. B. Haslam, R. Madan, Leptin receptor q223r polymorphism influences neutrophil mobilization after Clostridium difficile infection. Mucosal Immunol 11, 947–957 (2018).

32. C. M. Theriot, C. C. Koumpouras, P. E. Carlson, Bergin, II, D. M. Aronoff, V. B. Young, Cefoperazone-treated mice as an experimental platform to assess differential virulence of Clostridium difficile strains. Gut Microbes 2, 326–334 (2011).

33. A. J. McDermott, K. E. Higdon, R. Muraglia, J. R. Erb-Downward, N. R. Falkowski, R. A. McDonald, V. B. Young, G. B. Huffnagle, The role of Gr-1(+) cells and tumour necrosis factor-alpha signalling during Clostridium difficile colitis in mice. Immunology 144, 704–716 (2015).

34. L. L. Reber, C. M. Gillis, P. Starkl, F. Jönsson, R. Sibilano, T. Marichal, N. Gaudenzio, M. Bérard, S. Rogalla, C. H. Contag, P. Bruhns, S. J. Galli, Neutrophil myeloperoxidase diminishes the toxic effects and mortality induced by lipopolysaccharide. J Exp Med 214, 1249–1258 (2017).

35. G. S. Gullotta, D. De Feo, E. Friebel, A. Semerano, G. M. Scotti, A. Bergamaschi, E. Butti, E. Brambilla, A. Genchi, A. Capotondo, M. Gallizioli, S. Coviello, M. Piccoli, T. Vigo, P. Della Valle, P. Ronchi, G. Comi, A. D’Angelo, N. Maugeri, L. Roveri, A. Uccelli, B. Becher, G. Martino, M. Bacigaluppi, Age-induced alterations of granulopoiesis generate atypical neutrophils that aggravate stroke pathology. Nat Immunol 24, 925–940 (2023).

36. C. Trapnell, D. Cacchiarelli, J. Grimsby, P. Pokharel, S. Li, M. Morse, N. J. Lennon, K. J. Livak, T. S. Mikkelsen, J. L. Rinn, The dynamics and regulators of cell fate decisions are revealed by pseudotemporal ordering of single cells. Nat Biotechnol 32, 381–386 (2014).

37. X. Qiu, A. Hill, J. Packer, D. Lin, Y. A. Ma, C. Trapnell, Single-cell mRNA quantification and differential analysis with Census. Nat Methods 14, 309–315 (2017).

38. J. Cao, M. Spielmann, X. Qiu, X. Huang, D. M. Ibrahim, A. J. Hill, F. Zhang, S. Mundlos, L. Christiansen, F. J. Steemers, C. Trapnell, J. Shendure, The single-cell transcriptional landscape of mammalian organogenesis. Nature 566, 496–502 (2019).

39. J. M. Adrover, J. A. Nicolas-Avila, A. Hidalgo, Aging: A Temporal Dimension for Neutrophils. Trends Immunol 37, 334–345 (2016).

40. I. Miralda, S. M. Uriarte, K. R. McLeish, Multiple Phenotypic Changes Define Neutrophil Priming. Front Cell Infect Microbiol 7, 217 (2017).

41. J. Czepiel, G. Biesiada, T. Brzozowski, A. Ptak-Belowska, W. Perucki, M. Birczynska, A. Jurczyszyn, M. Strzalka, A. Targosz, A. Garlicki, The role of local and systemic cytokines in patients infected with Clostridium difficile. J Physiol Pharmacol 65, 695–703 (2014).

42. H. Yu, K. Chen, Y. Sun, M. Carter, K. W. Garey, T. C. Savidge, S. Devaraj, M. E. Tessier, E. C. von Rosenvinge, C. P. Kelly, M. F. Pasetti, H. Feng, Cytokines Are Markers of the Clostridium difficile-Induced Inflammatory Response and Predict Disease Severity. Clin Vaccine Immunol 24, (2017).

43. D. V. S. Costa, V. Moura-Neto, D. T. Bolick, R. L. Guerrant, J. A. Fawad, J. H. Shin, P. Medeiros, S. E. Ledwaba, G. L. Kolling, C. S. Martins, V. Venkataraman, C. A. Warren, G. A. C. Brito, S100B Inhibition Attenuates Intestinal Damage and Diarrhea Severity During Clostridioides difficile Infection by Modulating Inflammatory Response. Front Cell Infect Microbiol 11, 739874 (2021).

44. G. T. Nguyen, E. R. Green, J. Mecsas, Neutrophils to the ROScue: Mechanisms of NADPH Oxidase Activation and Bacterial Resistance. Front Cell Infect Microbiol 7, 373 (2017).

45. O. Soehnlein, S. Oehmcke, X. Ma, A. G. Rothfuchs, R. Frithiof, N. van Rooijen, M. Morgelin, H. Herwald, L. Lindbom, Neutrophil degranulation mediates severe lung damage triggered by streptococcal M1 protein. Eur Respir J 32, 405–412 (2008).

46. K. M. Oliveira-Costa, G. B. Menezes, H. A. Paula Neto, Neutrophil accumulation within tissues: A damage x healing dichotomy. Biomed Pharmacother 145, 112422 (2022).

47. M. Mittal, M. R. Siddiqui, K. Tran, S. P. Reddy, A. B. Malik, Reactive oxygen species in inflammation and tissue injury. Antioxid Redox Signal 20, 1126–1167 (2014).

48. K. R. Genschmer, D. W. Russell, C. Lal, T. Szul, P. E. Bratcher, B. D. Noerager, M. Abdul Roda, X. Xu, G. Rezonzew, L. Viera, B. S. Dobosh, C. Margaroli, T. H. Abdalla, R. W. King, C. M. McNicholas, J. M. Wells, M. T. Dransfield, R. Tirouvanziam, A. Gaggar, J. E. Blalock, Activated PMN Exosomes: Pathogenic Entities Causing Matrix Destruction and Disease in the Lung. Cell 176, 113–126 e115 (2019).

49. A. Othman, M. Sekheri, J. G. Filep, Roles of neutrophil granule proteins in orchestrating inflammation and immunity. FEBS J 289, 3932–3953 (2022).

50. L. Liu, B. Sun, Neutrophil pyroptosis: new perspectives on sepsis. Cell Mol Life Sci 76, 2031–2042 (2019).

51. E. Lefrancais, B. Mallavia, H. Zhuo, C. S. Calfee, M. R. Looney, Maladaptive role of neutrophil extracellular traps in pathogen-induced lung injury. JCI Insight 3, (2018).

52. J. C. Ryu, M. J. Kim, Y. Kwon, J. H. Oh, S. S. Yoon, S. J. Shin, J. H. Yoon, J. H. Ryu, Neutrophil pyroptosis mediates pathology of P. aeruginosa lung infection in the absence of the NADPH oxidase NOX2. Mucosal Immunol 10, 757–774 (2017).

53. A. Welin, F. Amirbeagi, K. Christenson, L. Bjorkman, H. Bjornsdottir, H. Forsman, C. Dahlgren, A. Karlsson, J. Bylund, The human neutrophil subsets defined by the presence or absence of OLFM4 both transmigrate into tissue in vivo and give rise to distinct NETs in vitro. PLoS One 8, e69575 (2013).

54. M. N. Alder, J. Mallela, A. M. Opoka, P. Lahni, D. A. Hildeman, H. R. Wong, Olfactomedin 4 marks a subset of neutrophils in mice. Innate Immun 25, 22–33 (2019).

55. W. Liu, G. P. Rodgers, Olfactomedin 4 Is a Biomarker for the Severity of Infectious Diseases. Open Forum Infect Dis 9, ofac061 (2022).

56. W. Liu, M. Yan, Y. Liu, R. Wang, C. Li, C. Deng, A. Singh, W. G. Coleman, Jr., G. P. Rodgers, Olfactomedin 4 down-regulates innate immunity against Helicobacter pylori infection. Proc Natl Acad Sci U S A 107, 11056–11061 (2010).

57. A. Banerjee, S. Shukla, A. D. Pandey, S. Goswami, B. Bandyopadhyay, V. Ramachandran, S. Das, A. Malhotra, A. Agarwal, S. Adhikari, M. Rahman, S. Chatterjee, N. Bhattacharya, N. Basu, P. Pandey, V. Sood, S. Vrati, RNA-Seq analysis of peripheral blood mononuclear cells reveals unique transcriptional signatures associated with disease progression in dengue patients. Transl Res 186, 62–78 e69 (2017).

58. H. K. Brand, I. M. Ahout, D. de Ridder, A. van Diepen, Y. Li, M. Zaalberg, A. Andeweg, N. Roeleveld, R. de Groot, A. Warris, P. W. Hermans, G. Ferwerda, F. J. Staal, Olfactomedin 4 Serves as a Marker for Disease Severity in Pediatric Respiratory Syncytial Virus (RSV) Infection. PLoS One 10, e0131927 (2015).

59. E. E. Mannick, J. R. Schurr, A. Zapata, J. J. Lentz, M. Gastanaduy, R. L. Cote, A. Delgado, P. Correa, H. Correa, Gene expression in gastric biopsies from patients infected with Helicobacter pylori. Scand J Gastroenterol 39, 1192–1200 (2004).

60. O. Ramilo, W. Allman, W. Chung, A. Mejias, M. Ardura, C. Glaser, K. M. Wittkowski, B. Piqueras, J. Banchereau, A. K. Palucka, D. Chaussabel, Gene expression patterns in blood leukocytes discriminate patients with acute infections. Blood 109, 2066–2077 (2007).

61. M. N. Alder, A. M. Opoka, P. Lahni, D. A. Hildeman, H. R. Wong, Olfactomedin-4 Is a Candidate Marker for a Pathogenic Neutrophil Subset in Septic Shock. Crit Care Med 45, e426–e432 (2017).

62. W. Liu, M. Yan, J. A. Sugui, H. Li, C. Xu, J. Joo, K. J. Kwon-Chung, W. G. Coleman, G. P. Rodgers, Olfm4 deletion enhances defense against Staphylococcus aureus in chronic granulomatous disease. J Clin Invest 123, 3751–3755 (2013).

63. L. C. McDonald, D. N. Gerding, S. Johnson, J. S. Bakken, K. C. Carroll, S. E. Coffin, E. R. Dubberke, K. W. Garey, C. V. Gould, C. Kelly, V. Loo, J. Shaklee Sammons, T. J. Sandora, M. H. Wilcox, Clinical Practice Guidelines for Clostridium difficile Infection in Adults and Children: 2017 Update by the Infectious Diseases Society of America (IDSA) and Society for Healthcare Epidemiology of America (SHEA). Clin Infect Dis 66, e1–e48 (2018).

64. P. D. Farooq, N. H. Urrunaga, D. M. Tang, E. C. von Rosenvinge, Pseudomembranous colitis. Dis Mon 61, 181–206 (2015).

65. R. C. Furze, S. M. Rankin, Neutrophil mobilization and clearance in the bone marrow. Immunology 125, 281–288 (2008).

66. K. N. Kangelaris, R. Clemens, X. Fang, A. Jauregui, T. Liu, K. Vessel, T. Deiss, P. Sinha, A. Leligdowicz, K. D. Liu, H. Zhuo, M. N. Alder, H. R. Wong, C. S. Calfee, C. Lowell, M. A. Matthay, A neutrophil subset defined by intracellular olfactomedin 4 is associated with mortality in sepsis. Am J Physiol Lung Cell Mol Physiol 320, L892–L902 (2021).

67. P. X. Liew, P. Kubes, The Neutrophil’s Role During Health and Disease. Physiol Rev 99, 1223–1248 (2019).

68. J. E. Stark, A. M. Opoka, J. Mallela, P. Devarajan, Q. Ma, N. C. Levinsky, K. F. Stringer, H. R. Wong, M. N. Alder, Juvenile OLFM4-null mice are protected from sepsis. Am J Physiol Renal Physiol 318, F809–F816 (2020).

69. D. J. Flanagan, T. J. Phesse, N. Barker, R. H. Schwab, N. Amin, J. Malaterre, D. E. Stange, C. J. Nowell, S. A. Currie, J. T. Saw, E. Beuchert, R. G. Ramsay, O. J. Sansom, M. Ernst, H. Clevers, E. Vincan, Frizzled7 functions as a Wnt receptor in intestinal epithelial Lgr5(+) stem cells. Stem Cell Reports 4, 759–767 (2015).

70. S. Jose, A. Mukherjee, O. Horrigan, K. D. R. Setchell, W. Zhang, M. E. Moreno-Fernandez, H. Andersen, D. Sharma, D. B. Haslam, S. Divanovic, R. Madan, Obeticholic acid ameliorates severity of Clostridioides difficile infection in high fat diet-induced obese mice. Mucosal Immunology, (2020).

71. A. B. Ryder, Y. Huang, H. Li, M. Zheng, X. Wang, C. W. Stratton, X. Xu, Y. W. Tang, Assessment of Clostridium difficile infections by quantitative detection of tcdB toxin by use of a real-time cell analysis system. J Clin Microbiol 48, 4129–4134 (2010).

72. O. Franzen, L. M. Gan, J. L. M. Bjorkegren, PanglaoDB: a web server for exploration of mouse and human single-cell RNA sequencing data. Database (Oxford) 2019, (2019).

73. S. Aibar, C. B. Gonzalez-Blas, T. Moerman, V. A. Huynh-Thu, H. Imrichova, G. Hulselmans, F. Rambow, J. C. Marine, P. Geurts, J. Aerts, J. van den Oord, Z. K. Atak, J. Wouters, S. Aerts, SCENIC: single-cell regulatory network inference and clustering. Nat Methods 14, 1083–1086 (2017).

74. D. Aran, A. P. Looney, L. Liu, E. Wu, V. Fong, A. Hsu, S. Chak, R. P. Naikawadi, P. J. Wolters, A. R. Abate, A. J. Butte, M. Bhattacharya, Reference-based analysis of lung single-cell sequencing reveals a transitional profibrotic macrophage. Nat Immunol 20, 163–172 (2019).

75. C. Tabula Muris, A single-cell transcriptomic atlas characterizes ageing tissues in the mouse. Nature 583, 590–595 (2020).

76. J. Chen, E. E. Bardes, B. J. Aronow, A. G. Jegga, ToppGene Suite for gene list enrichment analysis and candidate gene prioritization. Nucleic Acids Res 37, W305–311 (2009).

77. L. Zappia, A. Oshlack, Clustering trees: a visualization for evaluating clusterings at multiple resolutions. Gigascience 7, (2018).

78. G. Yu, L. G. Wang, Y. Han, Q. Y. He, clusterProfiler: an R package for comparing biological themes among gene clusters. OMICS 16, 284–287 (2012).

79. Y. Zhou, B. Zhou, L. Pache, M. Chang, A. H. Khodabakhshi, O. Tanaseichuk, C. Benner, S. K. Chanda, Metascape provides a biologist-oriented resource for the analysis of systems-level datasets. Nat Commun 10, 1523 (2019).

80. S. Nestorowa, F. K. Hamey, B. Pijuan Sala, E. Diamanti, M. Shepherd, E. Laurenti, N. K. Wilson, D. G. Kent, B. Gottgens, A single-cell resolution map of mouse hematopoietic stem and progenitor cell differentiation. Blood 128, e20–31 (2016).

