## Supplementary Figures for "Olfactomedin-4^+^ neutrophils exacerbate intestinal epithelial damage and worsen host survival after *Clostridioides difficile* infection"

Huber *et al.*

**The file includes:**

Supplementary Materials and Methods

Figs. S1 to S6

### Supplementary Materials and Methods

**Post-Sequencing Filtering and Quality Control**

Cell Ranger was used to generate a matrix file with expression counts for each sample, with genes as rows and cell Unique Molecular Identifier (UMI) as columns. HDF5 file formats were produced and utilized for analysis in Seurat using the Read10X_h5 function (*74*). In Seurat, data was merged and filtered. Cells below 1% nFeatures_RNA (poor quality cells) and above 92.3% (multiplet rate is 7.7% for 10,000 recovered cells) were excluded from analyses. Dead/dying cells were filtered out based on presence of >5% mitochondrial genes. Additionally, later in the analysis, cells with high hemoglobin genes (>5%) and high ribosomal genes (>3%) were filtered out. Following initial filtering, the standard Seurat analysis of integrated data was utilized. We split the merged object by infection status to create a list (SplitObject), normalized (NormalizeData) and identified variable features (FindVariableFeatures) for each dataset independently, selected variable features (SelectIntegrationFeatures) across datasets for integration, identified anchors (FindIntegrationAnchors), used anchors to integrate the datasets (IntegrateData), scaled the data (ScaleData), ran PCA (RunPCA) followed by UMAP (RunUMAP), found neighbors (FindNeighbors) at 1:24 dimensions, and lastly found clusters (FindClusters) at 0.5 resolution.

AUCell was utilized to identify cells in our dataset that are enriched in the genes from a neutrophil gene dataset obtained from PanglaoDB database. Cells enriched for these mouse neutrophil genes were then labeled as either “Neutrophil” or “non_Neutrophil”. Neutrophils obtained from AUCell annotation were used as the input for a second annotation process using SingleR. Neutrophils identified were compared to BM cells in Tabula Muris Senis dataset. Cells labeled “granulocyte”, “hematopoietic precursor cells”, and “granulocytopoietic cell” were retained for further analysis. Lastly, utilizing ToppGene database, clusters that possessed DEGs (using FindAllMarkers function) associated with other cell types were excluded from further analysis.

**Clustering Samples**

After filtering out non-neutrophils, uninfected and infected BM and blood samples were clustered at dim= 1:15 and at resolutions 0.2, 0.4, 0.8, and 1.2. Clustree analysis was performed to visualize how Seurat clusters changed at different resolutions, while BuildClusterTree function was used to create a hierarchical clustering dendrogram of transcriptomic similarities of neutrophil populations at each resolution. Further, FindAllMarkers was utilized to calculate differentially expressed genes (DEGs) amongst the neutrophil populations at each resolution. Based off data collected from Clustree analysis, BuildClusterTree, and DGE analysis, clusters present at resolutions 0.4 and 0.8 were utilized for final clusters for analysis.

### Trans epithelial electrical resistance (TEER) assay.

Target cells (were seeded after establishing background with medium alone) into the wells of 96-well E-Plates in 100µL of media. Cell growth was monitored with the xCELLigence RTCA system until they formed a monolayer (24 to 34 hours, depending on the experiment). Effector cells (sorted neutrophils; 100,000 cells/well) were then directly added to the wells. *C. difficile* toxins (25ng/mL) and lipopolysaccharide (1µg/mL) were added to epithelial-neutrophil cocultures to mimic *in vivo* CDI conditions. TEER measurements were captured every 15 minutes for up to 72 hours.

**Animal housing and clinical scoring**

To prevent cross-infection, infected mice were single-caged and were monitored every 12-24hrs. Each cage was provided with ad libitum feed, drinking water, and a secondary enrichment. Animals were monitored daily during the experiment to assess body weight loss and clinical disease. CDI severity was measured using a scoring system based on weight loss, coat ruffling, ocular discharge, activity level, posture, and diarrhea. Each parameter was scored on a scale from 0 to 3, with higher clinical scores indicative of more severe morbidity. Mice were euthanized if the score indicated intense morbidity (score >14) on any day of the experiment. Kaplan-Meier curves for overall survival were plotted for all four groups, and the statistical significance between groups was evaluated using the log-rank [Mantel-Cox] test.

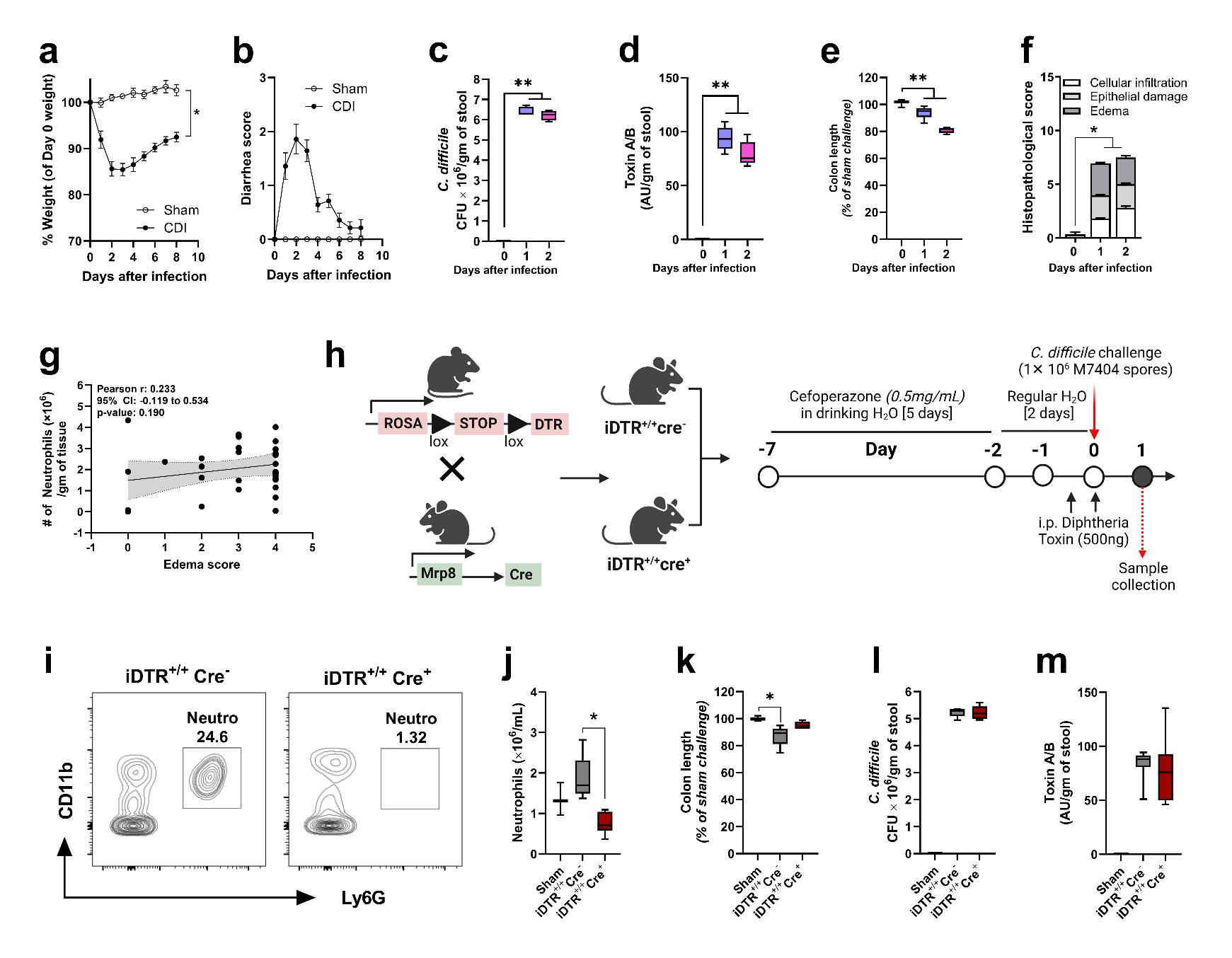

### Supplementary Figures

**Supplementary Figure 1: CDI clinical disease scores and pathogen burden in WT, iDTR^+/+^ Cre^-^ and iDTR^+/+^ Cre^+^ mice. a)** Percent weight loss **b)** diarrhea score of sham and *C. difficile*-challenged mice. **c)** *C. difficile* pathogen and **d)** toxin burden in cecal contents on day 0, day 1, and 2 of infection. **e)** colon length and **f)** histopathology score of *C. difficile* challenged C57 BL6 mice. **g)** Pearson correlation plot of tissue neutrophil number in the acute phase to cecal edema score. **h)** Schematic diagram of generation and *C. difficile* challenge of iDTR^+/+^ MRP8Cre^+^ mice. **i)** Representative FACS plot showing colonic neutrophils in iDTR^+/+^Cre^-^ (Control; top) and iDTR^+/+^Cre^+^ (bottom) mice on day 1 after infection. **j)** Number of neutrophils in blood collected on day 1 from sham or *C. difficile* challenged iDTR^+/+^Cre^-^ (control) and iDTR^+/+^Cre^+^ mice. **k)** colon length, **l)** pathogen burden **m)** and toxin titer in iDTR^+/+^Cre^-^ (control) and iDTR^+/+^Cre^+^ mice after *C. difficile* challenge. For panels a-f and j-m, data shown as mean ± SEM; N = 3-6 per group; representative of 2 independent experiments; *p < 0.05, Student’s t-test. For panel g, N = 33 pooled from multiple experiments.

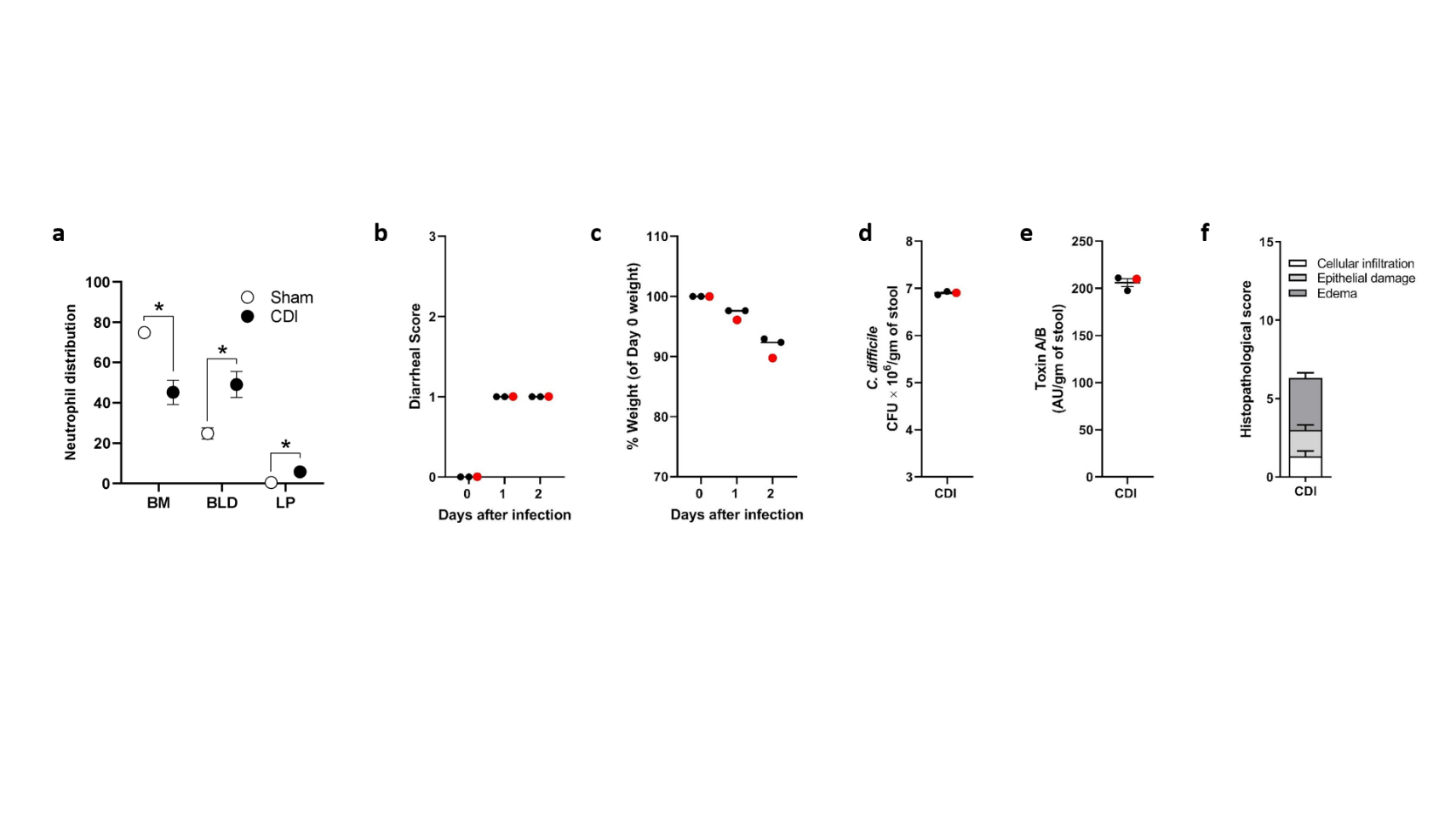

**Supplementary Figure 2: Neutrophil distributions after CDI and clinical disease score of *C. difficile*- infected mice used for neutrophil single-cell transcriptomics. a)** Distribution of neutrophils across bone marrow, blood, and colonic tissue of sham and *C. difficile*-challenged mice on day 2 after challenge. **b)** Diarrhea score, **c)** percent body weight, **d)** pathogen burden, **e)** cecal toxin level, and **f)** cecal histopathology score of C57BL/6 mice challenged with *C. difficile* spores for scRNA seq experiment. The mouse sample used for scRNA seq is highlighted in red. Panel a; N = 3-4 per group; representative of multiple experiments. Panel b-f; data shown as mean ± SEM; N = 3 per group; *p < 0.05, Student’s t-test.

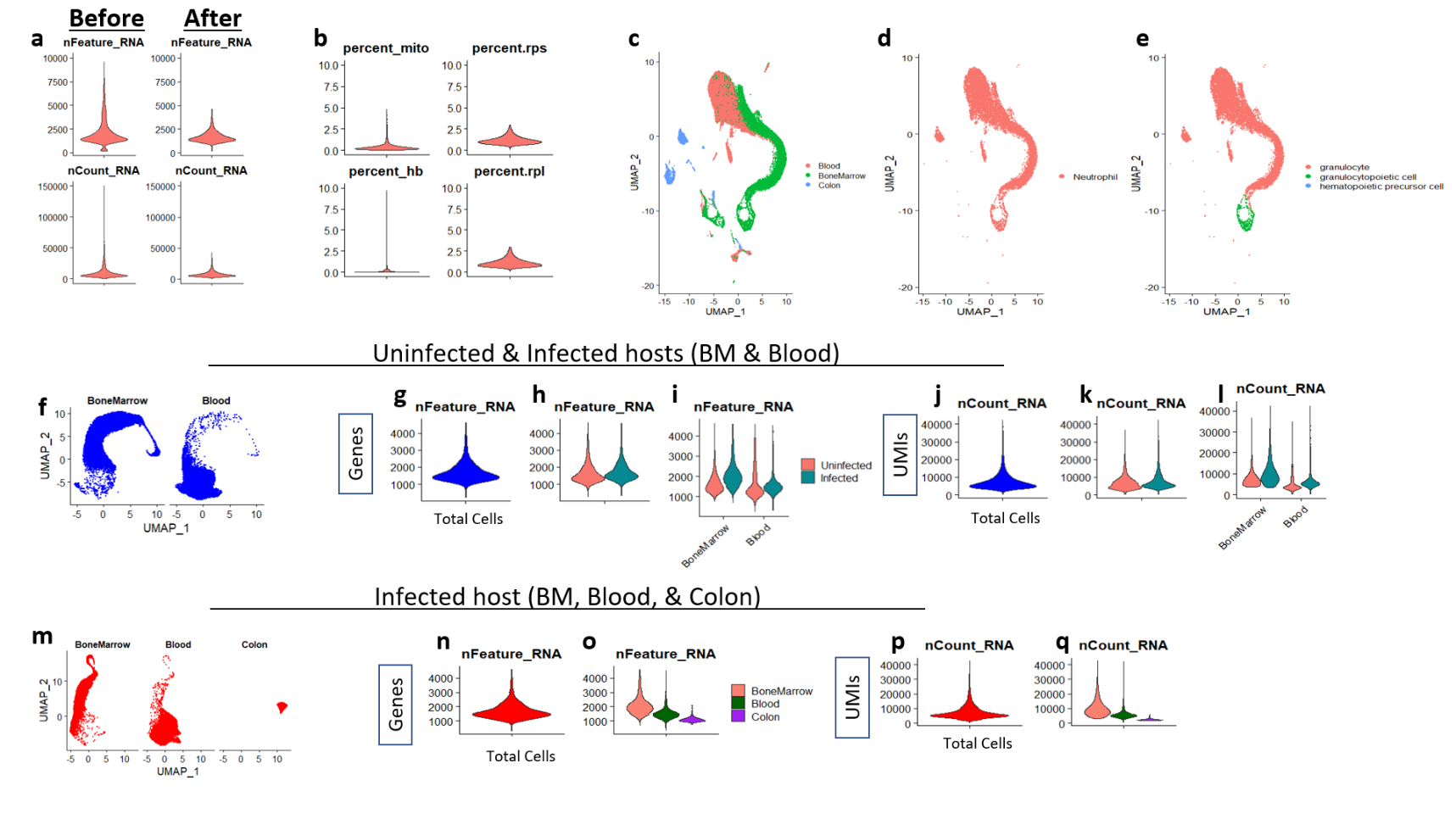

**Supplementary Figure 3: Quality control measures implemented in Sc-RNAseq analysis, identification of neutrophils, and Clustering. a)** Violin plots of cells before and after filtering out low quality cells (cells below 1% nFeatures_RNA) and doublets (cells above 92.3% nFeatures_RNA). **b)** Violin plots after filtering out cells with mitochondrial genes> 5%, hemoglobin genes>10%, and ribosomal genes>3%. **c)** UMAP of initial cell clustering. **d)** UMAP of cells enriched for neutrophil genes (termed “Neutrophil”). **e)** UMAP of cells labeled as cell types from the *Tabula Muris Senis* dataset. For initial analyses, sequences from BM and blood neutrophils of uninfected and infected host were integrated as one Seurat object. **f)** Violin plots of gene numbers per cell were graphed for the integrated Seurat object **g)** by total cells of the integrated object, **h)** by condition, and **i)** by condition and tissue. Violin plots of number of UMIs per cell and j**)** by total cells of the integrated object, **k)** by condition, and **l)** by condition and tissue. Number of genes per cell for the infected host Seurat object (BM, blood, and colon) plotted in violin plots **n)** together and **o)** by tissue. Number of UMIs per cell for the infected host Seurat objected plotted **p)** together and **q)** by tissue with violin plots.

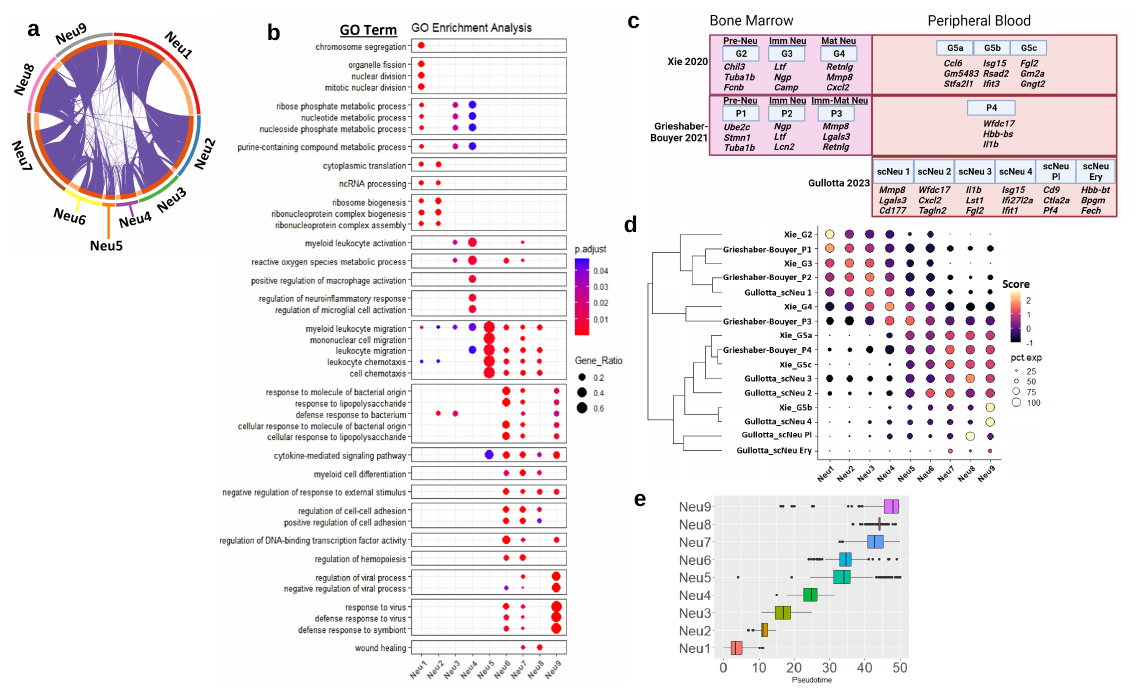

**Supplementary Figure 4**: **Characterization of neutrophil populations at steady state and during CDI and comparison with neutrophil populations defined in the literature.**  **a)** Circos plot of gene overlap analysis which displays shared DEGs among each cluster. **b)** Gene ontology analysis of neutrophil population DEGs were used to find biological functions enriched in each cluster. **c)** Signature DEGs of previously characterized mouse neutrophil populations in BM and blood at steady state (Xie 2020 and Grieshaber-Bouyer 2021), and in young/aged mice poststroke (Gullotta 2023). **d)** Scoring of neutrophil populations using DEGs of previously defined neutrophil populations. **e)** Mean pseudotime values for each cluster.

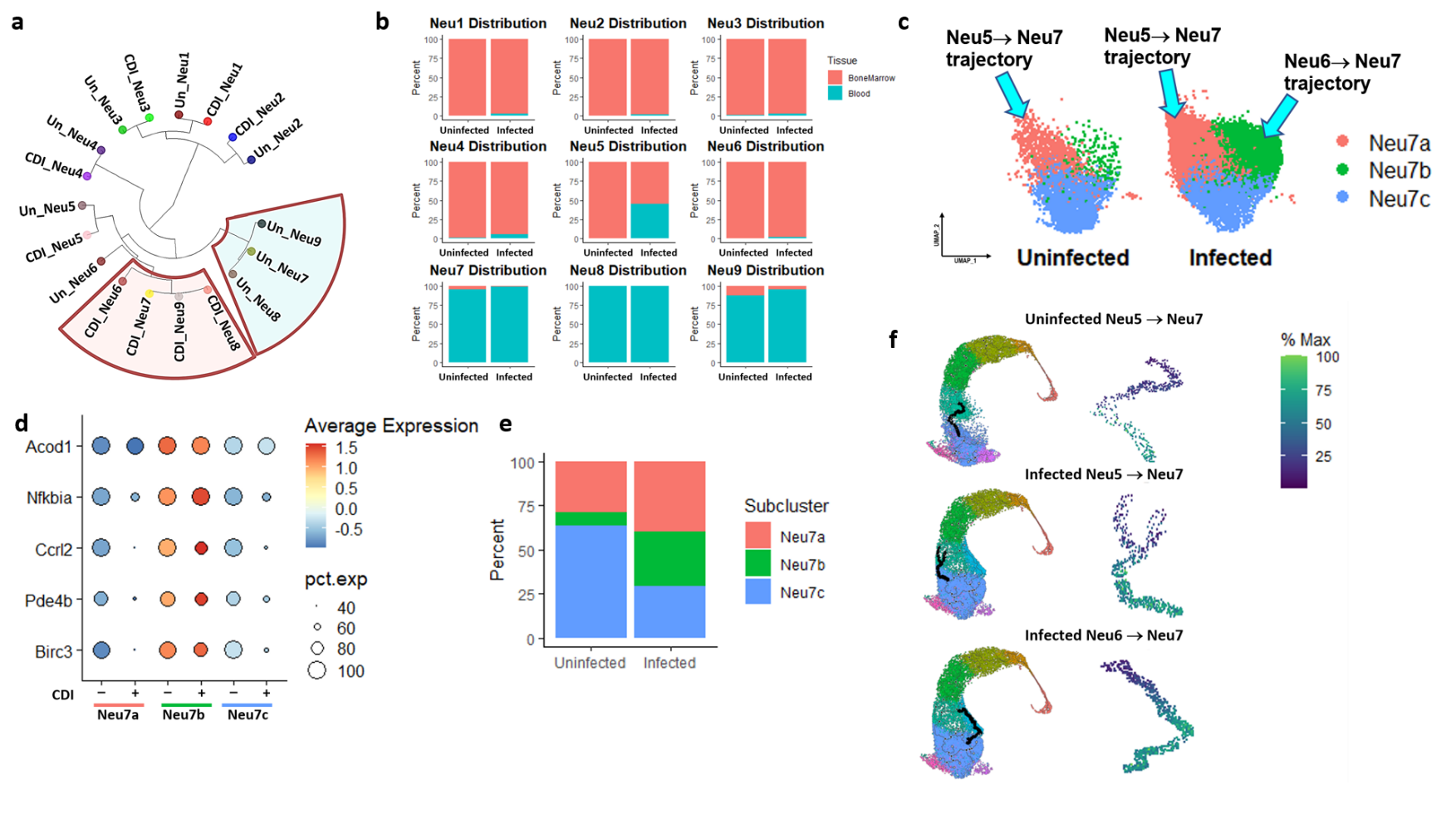
**Supplementary Figure 5: Characterization of *C. difficile*-induced changes to the neutrophil transcriptional landscape. a)** Cluster tree of neutrophil populations at steady state and during CDI. **b)**. Percent distribution of each neutrophil cluster by tissue location. **c)** UMAP depictions of Neu7 subpopulations of uninfected and *C. difficile*-infected hosts with locations in which differentiation trajectories give rise to Neu7a and Neu7b. **d)** Dot plot of DGE analysis of uninfected and infected host Neu7 subpopulations. **e)** Percent distribution of uninfected and infected host Neu7 subpopulations. **f)** Monocle branch analysis was utilized to calculate gene modules that change as a function of pseudotime along trajectories that give rise to Neu7.

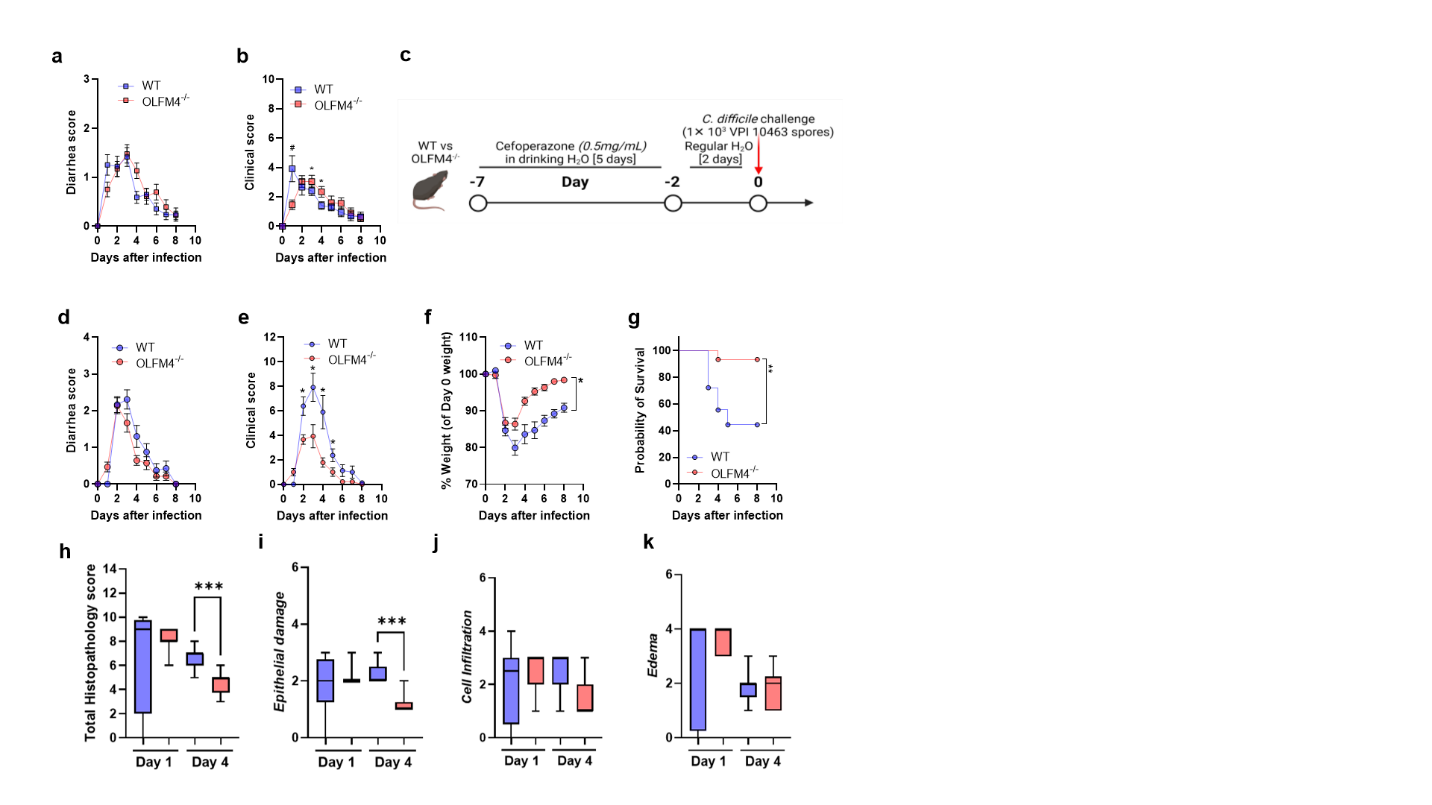

**Supplementary Figure 6: Clinical disease and histopathology scores of *C. difficile*-infected WT and OLFM4^-/-^ mice. a)** Diarrhea score and **b)** clinical disease score, after infection with M7404 spores. **c)** Schematic representation of the experimental design. Age- and gender-matched WT and OLFM4^-/-^ mice were pre-treated with antibiotics for 5 days in drinking water and challenged with 1× 10^3^ *C. difficile* (VPI 10463) spores by oro-gastric gavage two days after cessation of antibiotics. Animals were monitored until day 8 for survival. **d)** Diarrhea score, **e)** clinical disease score, **f)** percent weight change (compared to day 0 weight), and **g)** the difference in survival rate in mice challenged with VPI10463 spores (N = 15-18; Data shown as mean ± SEM; *p < 0.05; ANOVA or Log-rank [Mantel-Cox] test). **h)** Total histology score for **i)** epithelial damage, **j)** inflammatory cell infiltration, and **k)** submucosal edema in cecal tissue sections of *C. difficile*-challenged WT and OLFM4^-/-^ mice on day 1 and day 4 after challenge. N = 5-6; Data shown as mean ± SEM; N = 3 per group; representative of 2 independent experiments; *p < 0.05, **p < 0.01, ***p < 0.001; Student’s t-test.
